# Trogocytosis-mediated transfer of FOLR2 from nurse-like cells to CLL cells is associated with their activation and proliferation

**DOI:** 10.1101/2024.12.31.630890

**Authors:** Marcin Domagala, Chloe Bazile, Bastien Gerby, Loïc Ysebaert, Vera Pancaldi, Camille Laurent, Mary Poupot

**Author notes:** **Corresponding authors:** Marcin Domagala and Mary Poupot; 2 avenue Hubert Curien Oncopole de Toulouse, CS53717, 31037, Toulouse, France.

## Abstract

Communication with the lymphoid microenvironment is crucial for survival and proliferation of neoplastic B cells in chronic lymphocytic leukemia (CLL). In this study, we examined nurse-like cells (NLCs), CLL-specific macrophages, strongly implicated in CLL pathogenesis, and their interaction with leukemic cells. Using primary patient cells, we demonstrated that NLCs express high levels of folate receptor beta (FOLR2), which correlates with increased survival of cancer cells in vitro. Furthermore, we discovered that CLL cells acquire functional FOLR2 from NLCs via trogocytosis, enhancing their folate uptake. By mimicking the CLL microenvironment with soluble factors, CD40L and IL-15, we observed a strong NLC-dependent CLL cell activation and proliferation. Moreover, we linked this phenomenon with trogocytosis, by demonstrating that FOLR2+ CLL cells are the predominant population of actively cycling cancer cells. By multiplex immunofluorescence analysis of CLL patient lymph nodes, we confirmed the presence of FOLR2⁺ NLCs, and observed their enrichment in more aggressive and proliferative CLL cases. Finally, we detected FOLR2⁺ cancer cells, providing evidence of trogocytosis in situ. Taken together, we propose FOLR2 as a novel marker of protective NLCs, and highlight the importance of trogocytosis in improved CLL cell adaptation, including NLC-mediated activation and proliferation.

**Scheme 1.**
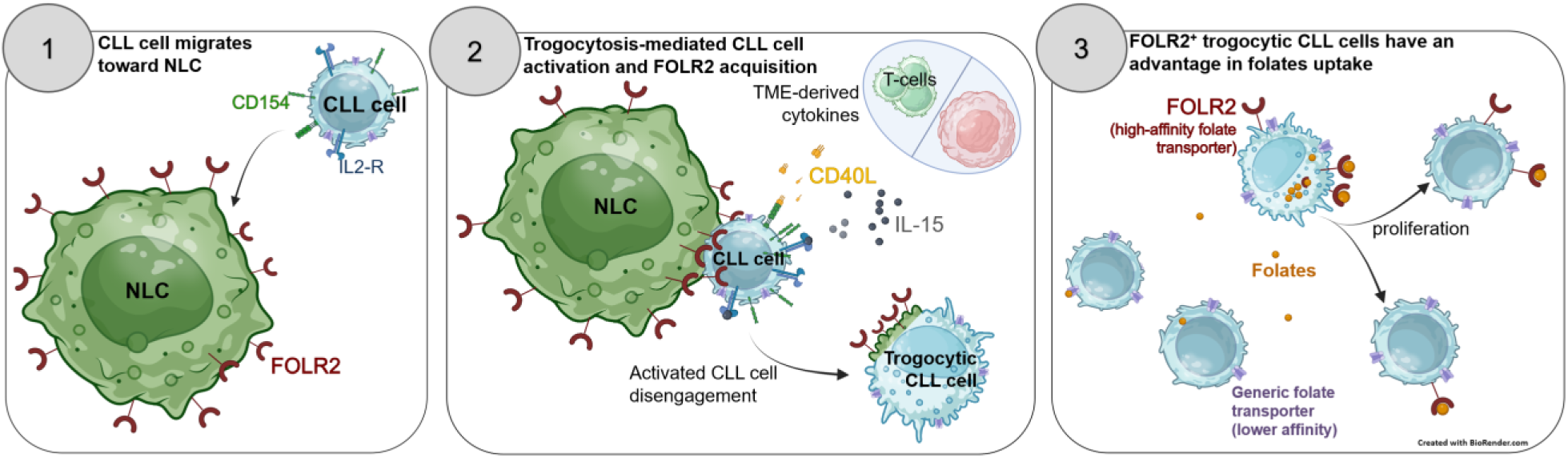
Schematic representation of trogocytosis-mediated acquisition of NLC-derived FOLR2 by CLL cells, and its impact on promoting cancer cell adaptability to folate deprivation. 1) CLL cells enter the lymph node (LN) and engage in contact with FOLR2^+^ NLCs. 2) The presence of T cell- and stromal cell-derived cytokines, CD40L and IL-15 promotes the activation and trogocytosis-mediated acquisition of FOLR2 by CLL cells. 3) FOLR2⁺ trogocytic CLL cells have an advantage in folate acquisition and show increased proliferation compared to the remaining population of cancer cells

**SUMMARY:**

- CLL cells acquire functional FOLR2 from NLCs via trogocytosis, a process that is closely linked with enhanced CLL cell activation and proliferation in vitro
- Increased Frequency of FOLR2 expressing macrophages in more aggressive and proliferative cases of CLL.
- FOLR2 is a marker of M2-like protective NLCs in vitro

## INTRODUCTION

Chronic lymphocytic leukemia (CLL) is the most common B cell malignancy in Western countries, characterized by the clonal accumulation of dysfunctional, CD5^+^ mature-like B cells in peripheral blood and lymphoid organs, including bone marrow and lymph nodes (LNs)^1^. CLL cells in the periphery are mostly quiescent and migrate to lymphoid organs, where they proliferate through interactions with the specific microenvironment^1^. Moreover, the CLL microenvironment, including: T cells, endothelial cells, stromal cells, and nurse-like cells (NLCs)^1,2^ is critical for cancer cell survival and chemoresistance^3–7^. NLCs were first observed in the *in vitro* CLL PBMC cultures ^8,9^, and described as a monocyte-derived population of macrophages, interacting closely with CLL cells, and protecting them against spontaneous apoptosis^8,9^, and therapeutic attacks^10–12^. Besides, NLCs are crucial for attracting cancer cells to the lymphoid organs^1^, and their increased numbers in the LNs and infiltration into proliferation centers correlates with more aggressive disease^5^ and shorter overall survival of the patients^5,13^.

The supportive role of NLCs toward CLL cells was demonstrated through the release of soluble factors^1,2,6,10,14^, extracellular vesicles^15^ as well as direct cell-cell interactions. Although numerous studies reported the clustering of CLL cells around NLCs^8,9,16^, only a few have further characterized the nature and functional impact of these direct interactions on cancer cells. For instance, vimentin and calreticulin presented by NLCs, induced activation of CLL cells, through B-cell receptors (BCRs), thereby promoting their survival and potential proliferation^2,17^. Moreover, direct contact between NLCs and CLL cells mediated through CD31 on NLCs and CD38/CD100 on CLL cells was implicated in the proliferation of CD38^+^/CD100^+^ malignant clones^18^. Furthermore, we showed that trogocytic interaction, partially mediated through CD2/CD58 (LFA-3), is essential for promoting cancer cell survival *in vitro*^19^. Trogocytosis, is a peculiar type of direct cell-cell interaction that leads to rapid acquisition of cellular content, including integral plasma membrane fragments^20^. This mechanism is predominantly unidirectional, enabling recipient cells, among others, to acquire new functionalities and modulate immune responses^20–22^. Nevertheless, the full impact of trogocytosis in CLL-NLC communication remains poorly understood, and requires further investigation.

Protumoral properties of NLCs are directly linked to their M2-like^12^, immunosuppressive polarization state^23,24^, typically defined by high expression of CD68^8,9^, CD163^5,12^. Indeed, treatment of NLCs with proinflammatory cytokines like IFNγ^25^, or TNF^26^, can abolish their nursing properties, including ability to interact with CLL cells, and induce M1-like polarization state, not always explained by CD163 expression. Here, we aimed to further characterize phenotype of NLCs, and their interactions with CLL cells. Using primary human samples, we observed that M2-like NLCs exhibit elevated levels of folate receptor beta (FOLR2), a myeloid cell-restricted^27^ high affinity transporter of folates^28^, which recently emerged as a specific marker of tumor-associated macrophages ^29–31^. We observed that FOLR2 expression on NLCs shows a markedly stronger association with *in vitro* CLL cell survival relative to CD163. Unexpectedly, we discovered that CLL cells acquire functional FOLR2 from NLCs via trogocytosis, and further defined the link between this interaction, and activation and proliferation of CLL cells. We supported our *in vitro* findings by detection of FOLR2^+^ NLCs in CLL patients LNs, their association with more aggressive, and proliferative disease, as well as discovery of FOLR2^+^ CLL cells *in situ*. With this study we propose FOLR2 as a novel marker of protumoral NLCs, and underline trogocytosis as an important mechanism of CLL-NLC communication, implicated in cancer cell activation and proliferation.

## MATERIALS AND METHODS

### Primary human cell isolation and culture

Peripheral blood samples from previously untreated CLL Patients were obtained from the Hematology Department with informed consent and referenced in the INSERM cell bank. According to French law, the INSERM cell bank has been registered with the Ministry of Higher Education and Research (DC-2013-1903) after being approved by an ethics committee (Comité de Protection des Personnes Sud-Ouest et Outremer II). Clinical and Biological annotations of the samples have been reported to the Comité National Informatique et Liberté. Buffy coats from healthy donors (HD) were obtained from Établissement français du sang (EFS, France).

Further details regarding cell isolation, *in vitro* culture conditions, and optimization of heterologous co-culture system, are listed in Supplementary Methods.

### Flow cytometry

Cells were stained with titrated antibodies (Supplementary Table S1) and analyzed immediately using LSR II or Fortessa X-20 flow cytometers (BD Pharmingen, France). Data were analyzed with FlowLogic 7.00.2 software (Inivai Technologies). A detailed description of staining protocols is provided in the Supplementary Methods. Gating strategies are presented in Supplementary Figures (S2A, S4E, S5A).

### FOLR2 gene knockout with RNP Crispr-Cas9

FOLR2 gene knockout (KO) in monocytes used for generation of HD-NLCs was achieved using ribonucleoprotein (RNP)-mediated CRISPR-Cas9 sgRNA method. Three predesigned crRNAs (IDT; sequences listed in Supplementary Table S1) targeting two alleles of FOLR2 gene or negative control sequences were delivered via electroporation Neon Transfection System (Thermo Fisher Scientific). Efficacy of FOLR2 KO was evaluated by qPCR (Fig. S2B) or flow cytometry (Fig. S2D). Further information is provided in Supplementary Methods.

### Trogocytosis assay

Autologous NLCs were detached and stained with PKH67 (Sigma) according to the manufacturer’s protocol. PKH67⁺ NLCs were then mixed with purified, naïve CLL cells at a 1:3 ratio in 96-well ultra-low attachment plates (Nunclon Sphera, Thermo Scientific), and incubated at 37°C for 4 h. Subsequently, cells were stained with appropriate antibodies and analyzed by flow cytometry. Purified naïve CLL cells cultured alone were used as a negative control of trogocytosis.

### Statistical analysis and results representation

Statistical significance and plotting of the results were performed using Graph Pad Prism 10.1.2 (USA). A paired, two-tailed t-test was used for comparison of two dependent groups, while repeated-measures ANOVA with Geisser–Greenhouse correction was employed for comparison of three or more dependent groups. Accordingly, an unpaired t-test and ordinary one-way ANOVA were used for comparisons between independent groups. Relation between CLL cell survival and specific marker expression on NLCs was calculated using Pearson correlation. The statistical test applied for each experiment is indicated in the respective figure legends. Results in the bar plots are presented as mean ± SD. The lines in the middle of the boxes in the box-and-whisker plots represent median values. The p-values below 0.05 were considered statistically significant.

Figures were edited with Inkscape software. Further information regarding experimental procedures, including: microscope imaging, and gene expression measurements, and result analysis are listed in Supplementary Methods. Details on key reagents, equipment and software employed in this study are provided in Supplementary Table S1.

## RESULTS

### FOLR2 expression by NLCs correlates with survival of CLL cells in vitro

The polarization state of TAMs is determinant for their role in the TME^32^. Previously, we showed that M2-like NLCs (CD163^high^, CD14^high^, CD209^pos^, CD64^low^, CD86^low^) interact with and support survival of CLL cells, unlike M1-like NLCs^26^. To further define the phenotype of protective NLCs, we used CLL PBMC *in vitro* cultures (Fig. 1A), and screened for protein markers associated with protumoral TAMs (Fig. S1A). This analysis revealed that M2-like NLCs display high levels of FOLR2 (Fig. S1A, 1B), further co-detected with CD163, a standard NLCs marker^5^ (Fig. S1B), and significantly overexpressed (MFI = 23057 vs 4268, n=9, p<0.003) compared with M2-like MDMs (Fig. 1C).

**Figure 1.**
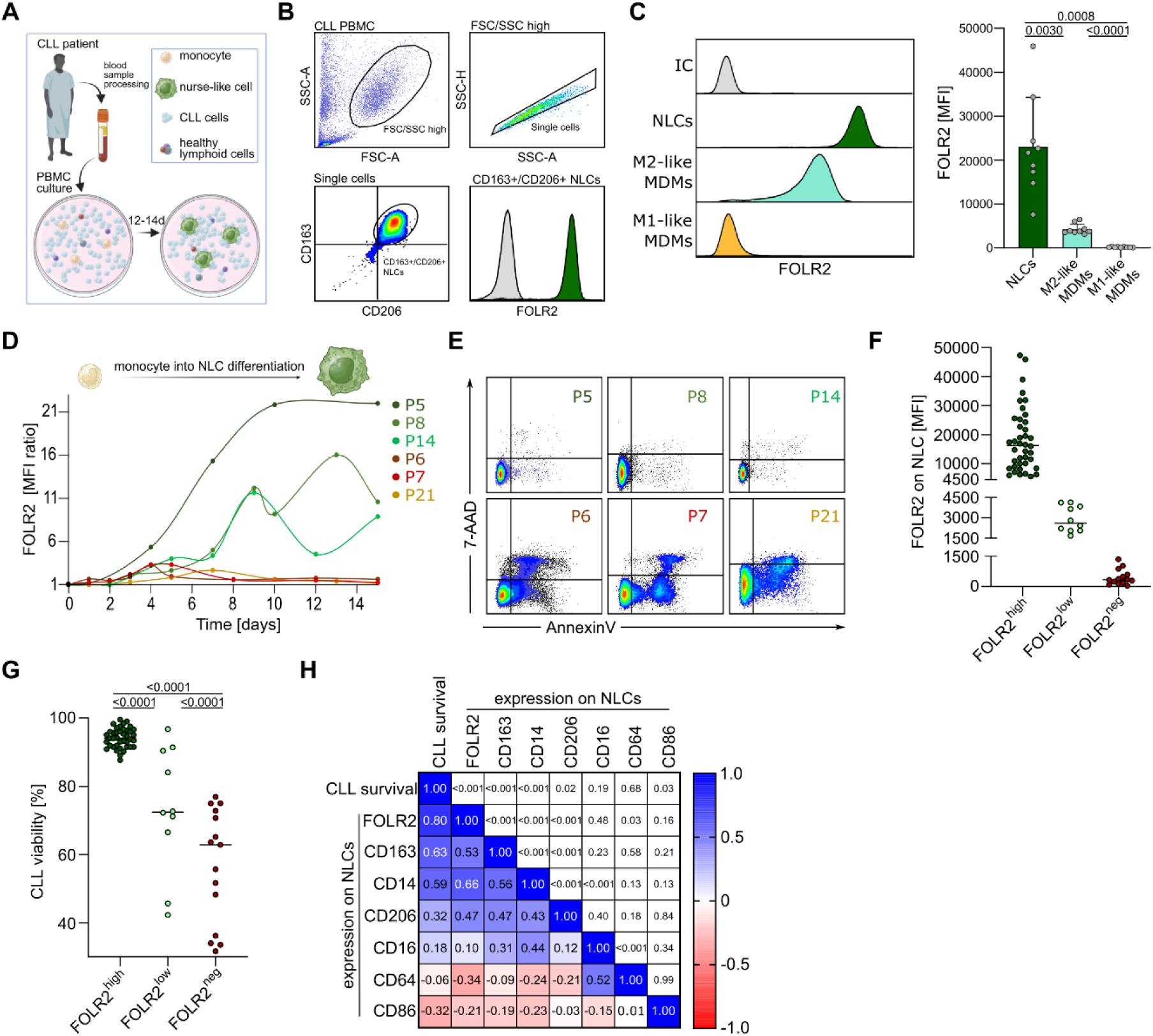
FOLR2 is a marker of protective NLCs. **A)** Schematic representation of NLC generation through high-density CLL PBMC cultures *in vitro*. **B)** Flow cytometry (FC) analysis of FOLR2 expression on a subpopulation of CD163⁺CD206⁺ NLCs (grey: isotype control; green: anti-FOLR2). **C)** Comparison of FOLR2 expression among different types of *in vitro*–generated macrophages: NLCs, M2-like, and M1-like monocyte-derived macrophages (MDMs); left: representative histograms; right: summary of results from n=9 independent experiments, each with cells from a different patient or donor. Statistical significance was evaluated with ordinary one-way ANOVA. **D)** FC measurement of FOLR2 expression by CD14^+^ cells during monocyte-to-NLC differentiation over 15 days of CLL PBMC culture (MFI ratio: anti-FOLR2/isotype control). Results for patients: P5-8, 14, 21 (numbers in brackets indicate initial percentage of monocytes in PBMC sample; each dot represents a measurement point). **E)** Dot plots with Annexin V/7-AAD staining of CLL cells from P5-8, 14, 21 measured on day 12 CLL PBMC culture. **F-G)** FC measurement of FOLR2 expression by NLCs and CLL viability for CLL PBMC cultures from n = 64 different patient samples (further details in the Supplementary Table 2), between day 12-14. **F)** Combined results of FOLR2 expression by NLCs, with samples stratified into three groups based on FOLR2 expression by NLCs and survival of CLL cells: FOLR2^neg^ (MFI < 1500, median = 340, n = 14); FOLR2^low^ (1500 < MFI < 4500, median = 3227, n = 10); FOLR2high (MFI>4500, median = 16258; CLL viability >85%; n = 40). **G)** Comparison of CLL survival in PBMC cultures, depending on FOLR2 expression by NLC, analyzed by ordinary one-way ANOVA. **H)** Analysis of Spearman correlation (r) between myeloid marker expression by NLCs and CLL cell viability between day 12-14 of the culture. The lower triangular of the matrix shows r values between tested parameters, according to color coded scale; the upper triangular of the matrix displays corresponding p-values. Results from n = 58 different patient samples. Results with p-values < 0.05 were considered statistically significant.

Next, we evaluated the kinetics of FOLR2 expression by CD14^+^ monocytes differentiating into NLCs over 15 days of PBMC culture. FOLR2 signal was detectable on CD14^+^ cells from day 3, peaking at days 9-10 for samples P5, P8, and P14 (FOLR2^+^ NLC), but diminishing completely toward the end of the culture for P6, P7, and P21 (FOLR2^-^ NLCs; Fig. 1D). Moreover, CLL cell survival exceeded 90% for the samples with FOLR2^+^ NLCs, but dropped below 75% in those with FOLR2^-^ NLCs (Fig. 1E). With systematic analysis of FOLR2 expression by NLCs and CLL survival across 64 patient samples (Fig. 1F-G; further details in Supplementary Table S2), we categorized PBMC cultures into three groups: FOLR2^high^ (MFI>4500, median=16258), FOLR2^low^ (1500<MFI<4500, median=3227) and FOLR2^neg^ (MFI<1500, median=340). CLL cell viability varied significantly among these groups (Fig.1G), being highest in the FOLR2^high^ (mean=94.2%, n=40) and the lowest in the FOLR2^neg^ (mean=57.5%, n=14). Furthermore, comparison between the PBMC and purified CLL cell cultures, showed significantly increased survival of cancer cells only in samples from FOLR2^high^ group (Fig. S1C). Finally, among analyzed myeloid markers on NLCs, expression of FOLR2 showed the strongest correlation with survival of CLL cells (Fig. 1H), followed by CD163 and CD14, whereas the M1 markers CD64 and CD86 displayed a negative association with cancer cell survival.

These results indicate FOLR2 as a specific marker of protective M2-like NLCs that support CLL cell survival *in vitro*.

### FOLR2 is detected on CLL cells co-cultured with NLCs

Given that CLL cells drive the maturation of FOLR2^+^ NLCs, which in turn protect cancer cells from apoptosis^13,33^, we explored the potential connection between FOLR2 and the interaction between these two cell types.

During standard phenotyping of NLCs, we observed a subpopulation of lymphoid cells with varying levels of myeloid markers: CD14, CD163, CD206 and FOLR2 (Fig. 2A; Fig.S2A for gating strategy), among which only FOLR2 was significantly detected (Fig.2B). Further analyses confirmed that FOLR2 is absent on freshly isolated CD5^+^/CD19^+^ CLL cells, and appears only after long-term PBMC culture (Fig. 2C), despite the lack of *FOLR2* gene expression by cancer cells (Fig. S2B). These results suggested that CLL cells acquire FOLR2 from other cells present in PBMC cultures, most likely NLCs.

**Figure 2.**
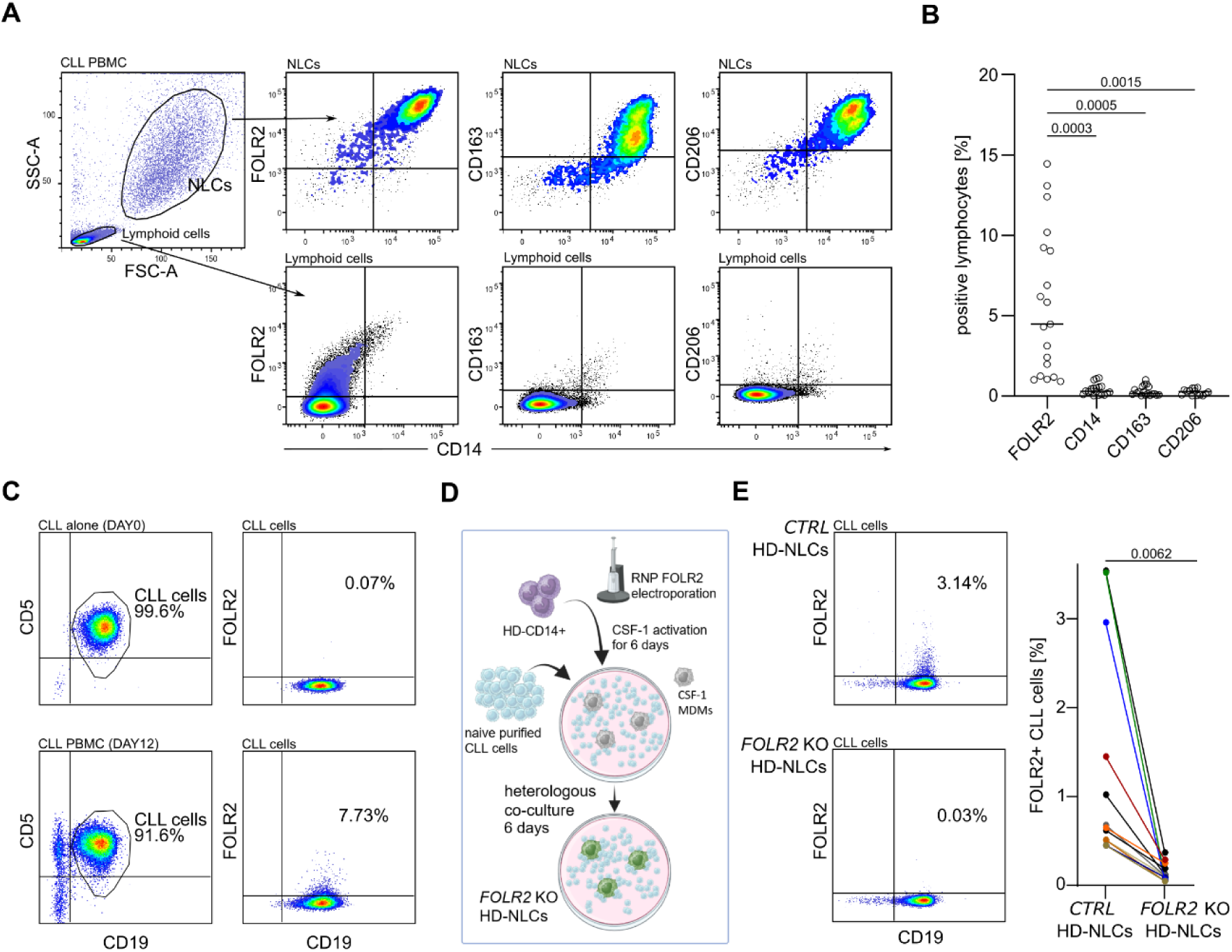
Detection of FOLR2 on the surface of CLL cells. **A)** Representative dot plots from flow cytometry (FC) analysis of FOLR2, CD14, CD163, and CD206 expression on cells from CLL PBMC cultures between day 12-14. Upper panels: results for large, granular cells corresponding to NLCs; lower panels: results for small cells corresponding to lymphoid cells. **B)** Frequency of FOLR2, CD14, CD163, and CD206 positive lymphoid cells (including CLL cells) from CLL PBMC cultures at day 12; results from n = 19 different patient samples. Statistical comparison was performed with repeated-measures ANOVA. **C)** FC analysis of FOLR2 expression on purified CLL cells (CD5⁺/CD19⁺) right after isolation from blood (day0) or after 12 days of PBMC culture. **D)** Schematic representation of FOLR2 knockout (KO) generation in NLCs using a ribonucleoprotein (RNP) CRISPR–Cas9 approach, combined with heterologous co-culture system composed of healthy donor (HD) CD14⁺ monocytes and purified CLL cells. **E)** Effect of *FOLR2* KO in NLCs on FOLR2 detection on CD5⁺CD19⁺ CLL cells in heterologous co-cultures: left – exemplary dot plots showing FOLR2 signal on CLL cells after co-culture with control (CTRL) HD-NLCs or *FOLR2* KO HD-NLCs; right: combined results of FOLR2⁺ CLL cells in co-cultures with CTRL HD-NLCs or *FOLR2* KO HD-NLCs (n = 5 independent experiments; 4 different patient samples; 12 measurements in total). Results with p-values < 0.05 were considered statistically significant.

To verify this hypothesis, we performed *FOLR2* knockout (KO) in NLCs. To generate sufficient NLC number for KO experiments, we used healthy donor NLC (HD-NLC) system, where HD-MDMs are co-cultured with purified CLL cells (Fig. 2D). HD-NLCs correspond to NLCs in terms of functionality^9^ and comparable FOLR2 expression (Fig. S2C). Co-cultures with *FOLR2* KO HD-NLCs led to a significant decrease in the percentage of FOLR2⁺ CLL cells compared with the control condition (Fig. 2E), confirming that NLCs are the source of FOLR2 detected on CLL cells.

### CLL cells acquire functional FOLR2 from NLCs through trogocytosis

Since CLL cells were shown to engage in trogocytosis with NLCs^19^, we investigated whether this mechanism is responsible for FOLR2 acquisition by cancer cells.

Following 4 hours co-culture with PKH-67 stained NLCs, we observed subpopulation of PKH-67+ and FOLR2+ CLL cells (Fig. 3A). Across 15 measurements, a mean of 20% and 2% of CLL cells were positive for PKH67 and FOLR2 respectively, while the signals from CD14 and CD16 were absent (Fig. 3B). Next, with confocal microscopy, we demonstrated direct interaction between FOLR2+ NLCs and CD19+ CLL cells (Fig. 3C), and observed FOLR2 patches co-localized with plasma membranes of cancer cells (Fig. 3D and S3A), which was abolished in co-cultures with FOLR2-KO HD-NLCs (Fig. 3E). These results indicate that CLL cells acquire FOLR2 via trogocytosis, and that the protein transfer is partially selective.

**Figure 3.**
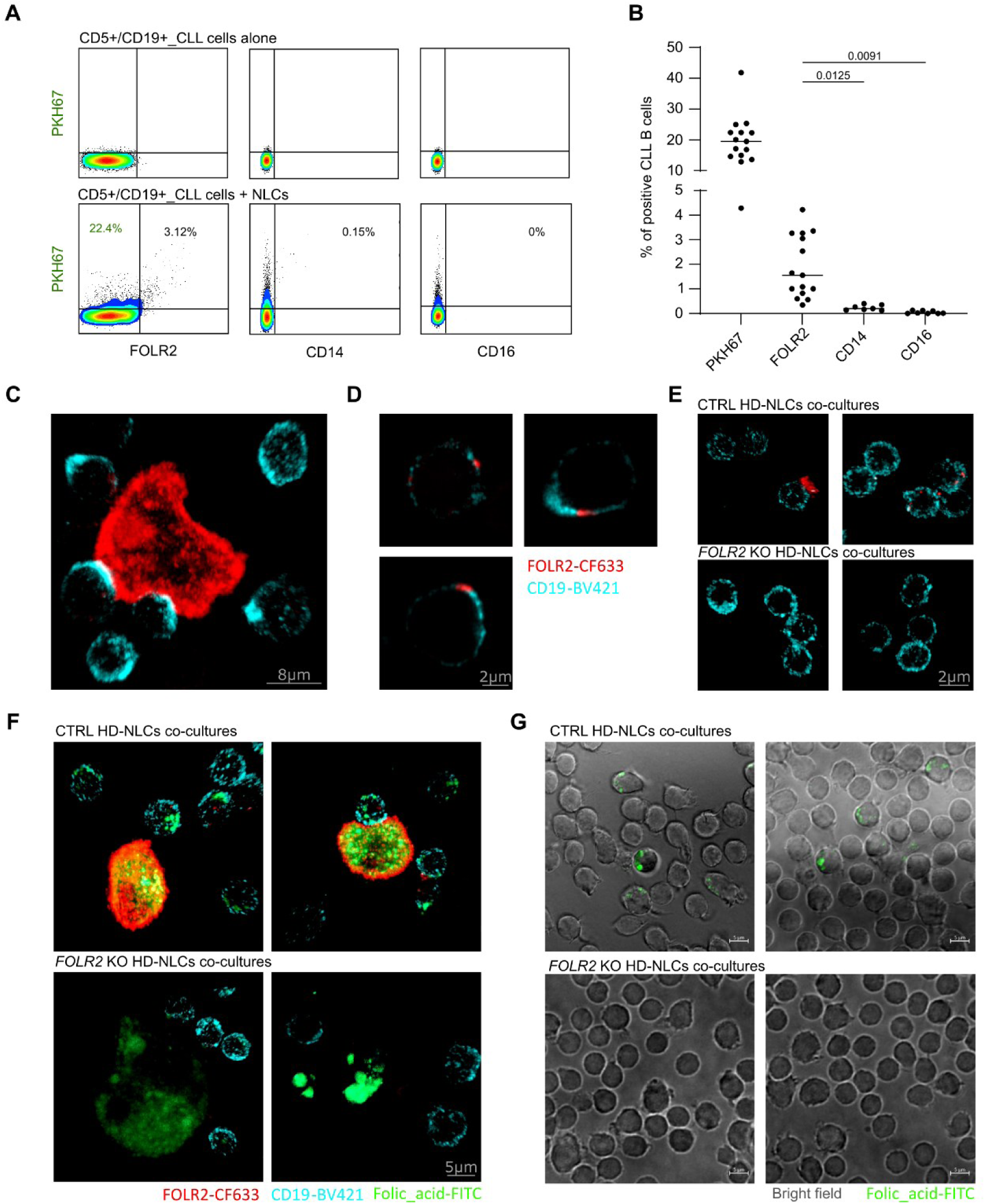
CLL cells acquire functional FOLR2 from NLCs by trogocytosis. **A)** Representative dot plots showing the percentage of CLL cells positive for PKH67, FOLR2, CD14, and CD16 after 4 hours of co-culture with PKH67⁺ NLCs (upper: CLL cells alone; lower: co-culture with NLCs). **B**) Combination of results from n = 3 independent experiments (cells from 5 patients; 15 measurements for PKH67 and FOLR2, 7 for CD14, and 8 for CD16). Statistical comparison between percentage of CLL cells positive for FOLR2, CD14 and CD16 was calculated by repeated-measures ANOVA. Results with p-values < 0.05 were considered statistically significant. **C–F)** Confocal microscopy images of CLL–NLC co-cultures stained with anti-CD19 (cyan) and anti-FOLR2 (red) antibodies: **C)** cell–cell contact between NLCs and CLL cells, represented as 3D projection; **D)** detection of FOLR2 patches on CLL cells following contact with NLCs; **E)** absence of FOLR2 signal on CLL cells after co-culture with *FOLR2* KO HD-NLCs; images represented as 3D projections. **F)** Confocal microscopy of CLL co-cultures with control (CTRL) HD-NLCs (upper) or FOLR2 KO HD-NLCs (lower), incubated with folic acid–FITC (green) for 90 min; images represented as 3D projections **G)** Epifluorescent images of purified CLL cells after co-culture with CTRL HD-NLCs (upper) or FOLR2 KO HD-NLCs (lower), followed by 90min incubation with folic acid–FITC.

To test whether the FOLR2 acquired by CLL cells is functional, we incubated the co-cultures with EC17, a folic acid-FITC imaging agent^32^. Expectedly, FOLR2^+^ NLCs showed higher intracellular EC17 signals, compared to FOLR2 KO NLCs (Fig. 3F). As for CD19^+^ CLL cells, we observed some with FOLR2 positive membrane patches and intracellular EC17 signal (Fig. 3F, top), which was greatly reduced in FOLR2 KO conditions (Fig. 3F, bottom). Moreover, only CLL cells purified after the co-culture with CTRL HD-NLCs, but not FOLR2 KO HD-NLCs were positive for EC17 (Fig. 3G). Finally, using flow cytometry, we observed an increased signal from the folic acid fluorescent probe in the subpopulation of FOLR2high CLL cells, as compared to FOLR2neg CLL cells (Fig. S3B). The data indicate that FOLR2 acquired via trogocytosis is functional, facilitating uptake of folates by CLL cells.

### Trogocytic FOLR2^+^ CLL cells are predominantly activated

Next, we tested whether FOLR2 acquisition by CLL cells is linked to their activation and proliferation. Considering that in patients, CLL cells could engage in trogocytosis only in the presence of TME, further required for their proliferation, we aimed to optimize culture conditions to mimic LN microenvironment. To generate sufficient numbers of macrophages observed in lymphoid organs, we used HD-NLC co-culture system, and substituted other cellular components of TME with soluble factors (Fig. S4A).

First, we compared IL-2+CpG^34^ and CD40L^1^+IL-15^34–36^ treatments to achieve more TLR- versus BCR-centric activation, respectively. We selected CD40L+IL-15 stimulation, which induced NLC-dependent CLL cell activation, as measured by CD23^37^ upregulation and CD184^38^ downregulation (Fig. S4B_A), and also increased the frequency of FOLR2⁺ CLL cells (fig. S4B_B). Next, we determined that an initial 1:9 proportion of macrophages to cancer cells best supports CLL–NLC trogocytosis, as measured by the increased frequency of FOLR2⁺ CLL cells (Fig. S4C). We finalized the activation protocol by adding cytokine treatment and medium exchange to minimize nutrient depletion (Fig. 4A).

**Figure 4.**
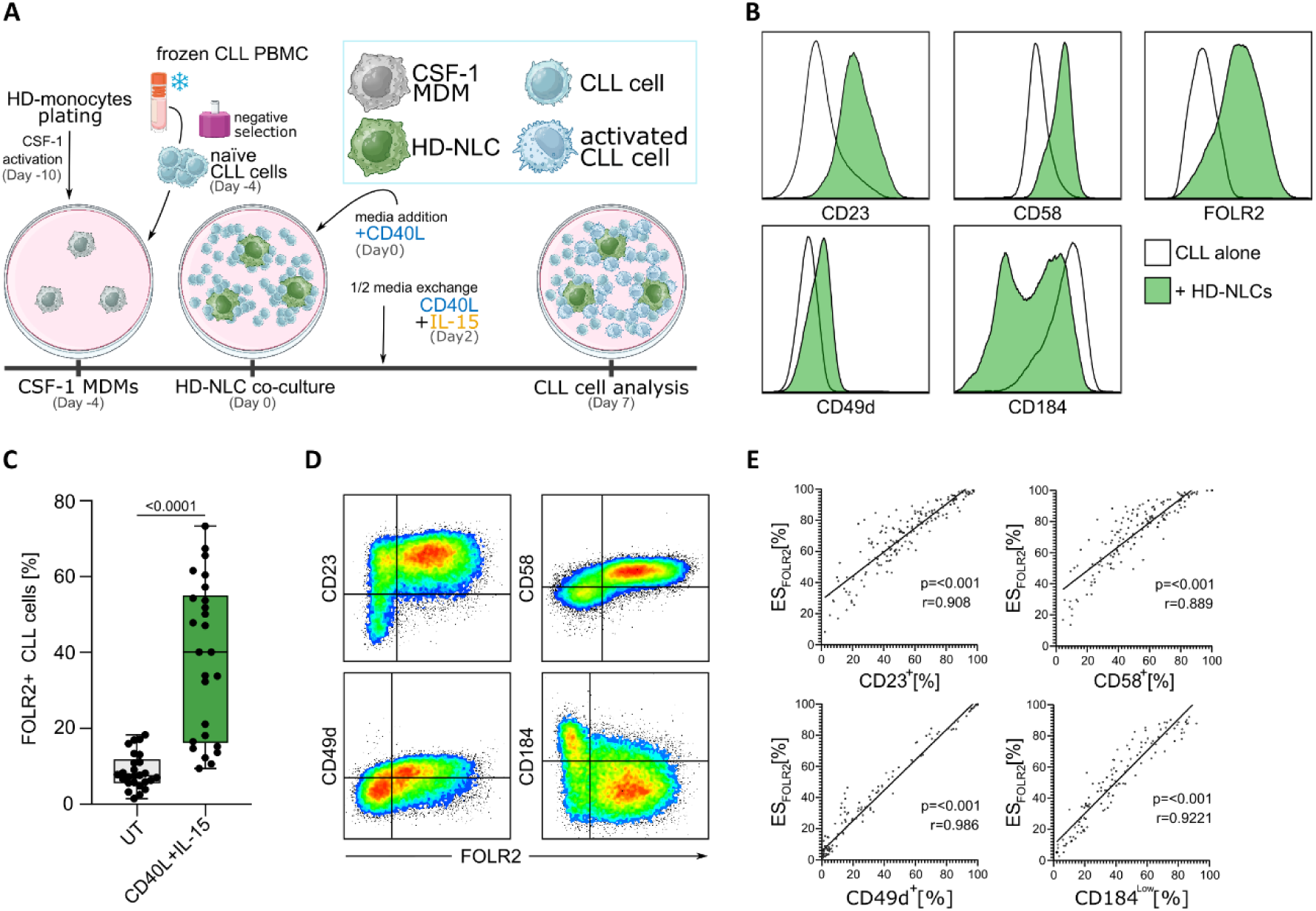
FOLR2 acquisition by CLL cells is positively associated with their activation. **A)** Schematic representation of optimized heterologous co-culture system for studying impact of trogocytosis on CLL cell activation and proliferation. **B)** Representative histograms from the flow cytometry (FC) analysis of CD23, CD58, CD49d and CD184 activation markers and FOLR2 levels on CD40L+IL-15 activated CLL cells, cultured alone (white) or in the presence of HD-NLCs (green). **C)** Comparison of FOLR2^+^ CLL cells percentage, following co-culture with HD-NLCs, in the presence or not of CD40L+IL-15. Statistical significance was determined by paired t-test. **D)** Dot plot representing co-expression of FOLR2 and CD23/ CD58/CD49d/CD184 on CLL cells after co-culture with HD-NLCs in the presence of CD40+IL-15. **E)** Pearson correlation between the total fraction of CLL cells expressing CD23/CD58/CD49d (positive activation markers) or CD184 (negative activation marker), and FOLR2 enrichment score (ES_FOLR2_; Fig.S4F_A), indicating proportion of the FOLR2^+^ cells detected in the population of activated cells (n=5 independent experiments, cells from 12 different patients; 150 measurement points). Results with p-values < 0.05 were considered statistically significant.

Using the optimized protocol, we confirmed significantly increased cancer cell survival in co-cultures with HD-NLCs compared to CLL cells cultured alone (Fig. S4D). Moreover, we observed increased expression of positive activation markers: CD23, CD49d^39,40^, CD58^18,42^, and decreased expression of negative activation marker CD184 by CLL cells co-cultured with HD-NLCs, relative to monocultures (Fig. 4B). Next, we demonstrated a significantly higher proportion of FOLR2^+^ CLL cells upon CD40L+IL-15 activation (9 vs 40%; Fig. 4C), which further led to detection of myeloid markers CD11b and CD14 on cancer cells (Fig. S4G).

Subsequently, we compared activation status of FOLR2^-^ and FOLR2^+^ CLL cells, and observed a higher proportion of FOLR2^+^ cells in the CD23^+^, CD49d^+^, CD58^+^, CD184^low^ subpopulation of cancer cells (Fig. 4D). We further supported these findings by demonstrating significant correlation between FOLR2 enrichment score (ES_FORL2_; Fig.S4F_A) and the total percentage of CLL cells expressing the corresponding activation marker (Fig. 4E).

Taken together, these results demonstrate that CD40L+IL-15 treatment induces strong NLC-dependent activation of cancer cells, greatly promotes acquisition of FOLR2 by CLL cells, and that FOLR2+ trogocytic CLL cells are predominant population of activated cancer cells.

### FOLR2^+^ CLL cells are predominant subpopulation of proliferating cancer cells

Next, we explored the potential impact of FOLR2 acquisition on CLL cell proliferation. First, we tested the impact of HD-NLCs and CD40L+IL-15 activation on Ki67 expression by CLL cells. We observed that the presence of HD-NLCs leads to robust induction of Ki67 expression by CLL cells (Fig. 5A, Fig. S5A for gating strategy), that depends on CD40L+IL-15 activation (Fig. S5B). Since Ki67 indicates actively cycling cells^43^, we performed dye-dilution assay using CellTrace™ Violet (CTV), to capture total fraction of cells after division, which further confirmed that co-culture with HD-NLCs induces significant proliferation of CLL cells upon CD40L+IL-15 stimulation (Fig. S5C).

**Figure 5.**
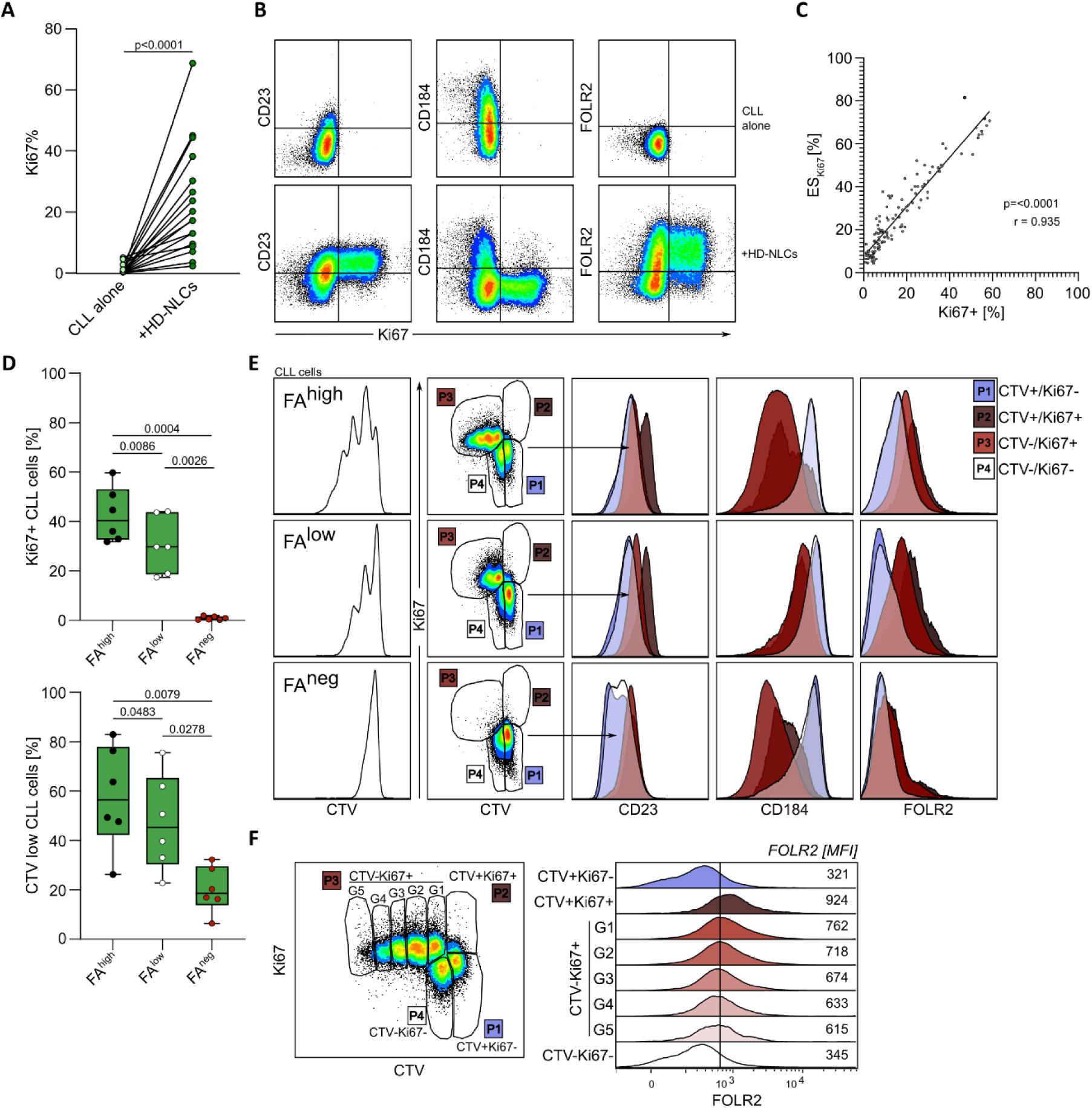
Trogocytic FOLR2^+^ CLL cells are predominant population of proliferating cancer cells. **A)** CLL monocultures or HD-NLCs co-cultures were activated with CD40L+IL-15, and Ki67 expression by CLL cells was measured by flow cytometry (FC). Comparison of the percentage of Ki67^+^ CLL cells between two culture modes, was assessed with paired t-test (n = 3 independent experiments; samples from 12 patients; 17 measurement points). **B)** Representative dot plots from FC analysis showing Ki67, CD23, CD184, and FOLR2 levels in CLL cells treated with CD40L+IL-15, and cultured either alone or with HD-NLCs. **C)** Pearson correlation between the total fraction of Ki67⁺ CLL cells and the Ki67 enrichment score (ESₖᵢ₆₇, Fig. S4F_B), representing the proportion of Ki67⁺ cells within the FOLR2⁺ trogocytic CLL cell population (n = 5 independent experiments; samples from 12 patients; 150 measurement points). **D)** Percentage of actively cycling (Ki67⁺, left) and total cycling (CTV^low^, right) fraction of CLL cells after co-culture with HD-NLCs, in the presence of CD40L+IL-15, under varying concentrations of folic acid (FA); cells from n = 6 different patient samples. Significance of the results was evaluated with repeated-measures ANOVA test with Geisser-Greenhouse correction. **E)** FC analysis of CD23, CD184, and FOLR2 signals on CLL cells gated based on Ki67/CTV levels: P1 = CTV^high^/Ki67⁻ (quiescent, non-dividing); P2 = CTV^high^/Ki67⁺ (right before first division); P3 = CTV^low^/Ki67⁺ (actively cycling, after division); P4 = CTV^low^/Ki67⁻ (post-cycling, quiescent). **F)** Exemplary stacked histogram of FOLR2 expression (MFI – median fluorescence intensity) on CLL cells according to their proliferative status (Ki67/CTV) and number of cell generations (G1–G5). Results with p-values < 0.05 were considered statistically significant.

Afterward, we examined the relationship between FOLR2 levels and proliferation status of CLL cells. We detected the majority of Ki67+ CLL cells in the activated CD23+/CD184low and FOLR2+ trogocytic subpopulations of cancer cells (Fig.5B). By calculating a Ki67 enrichment score (ES_Ki67_; Fig. S4F), representing the proportion of Ki67⁺ events within the FOLR2⁺ cell fraction, we found a strong correlation between ES_Ki67_ and total percentage of Ki67^+^ CLL cells (Fig. 5C), demonstrating FOLR2+ CLL cell enrichment in the fraction of actively proliferating cells.

Subsequently, we investigated the impact of folic acid (FA) availability on FOLR2 levels on CLL cells. First, we checked an effect of folic acid deprivation on CLL cells proliferation by testing three culture conditions with decreasing FA content: FAhigh (>1000 µg/L), FAlow (∼1.22 µg/L) and FAnegative (∼0 µg/L), using patient samples that previously showed high proliferation. We observed that FA reduction limits CLL proliferation in a dose-dependent manner, as indicated by significantly decreased Ki67 expression (Fig. 5D, top) and CTV dilution (Fig. 5D, bottom). Moreover, depletion of FA led to significantly smaller proportions of FOLR2 levels on CLL cells (Fig. S5D). Subsequently, we studied distribution of FOLR2+ CLL cells in the subpopulation of actively cycling cells, depending on the availability of FA (Fig. 5E). By combining Ki67 and CTV staining, we defined 4 subpopulations of CLL cells: P1. CTVhighKi67-, quiescent cells; P2. CTVhighKi67+, cells right before the first division; P3. CTVlowKi67+, subsequent generations of actively cycling cells; P4. CTVlowKi67-, cells after cycling, quiescent again (Fig. 5E). Analysis of FOLR2 levels in each subpopulation, revealed its dynamic fluctuations, with low levels before the first division (P1), significant increase for actively proliferating cells (P2, P3), and significant decrease in cells that had ceased proliferating (P4; Fig. S5E), irrespectively of initial FA content. Finally, we observed stable levels of FOLR2 in the subsequent generations of actively proliferating cells (Fig. 5F), indicating repeated engagement in trogocytosis by CLL cells between each cell division.

These experiments showed that FOLR2^+^ CLL cells are the predominant subpopulation of actively cycling CLL cells, and that trogocytic interaction with HD-NLCs resulting in FOLR2 acquisition is crucial for NLC-dependent proliferation of CLL cells upon CD40L+IL-15 treatment.

### Detection of trogocytic FOLR2+ CLL cells in the lymph nodes of CLL patients

Finally, we studied FOLR2 expression in tissues from CLL patients (supplementary table S3), to find a potential link with our in vitro findings. Using multiplex immunofluorescence (mIF), we measured FOLR2 and CD163 by macrophages in the reactive tonsil (n=1) from healthy donor, and LNs from patients with CLL cases ranging from classical CLL (cCLL; n=5), to accelerated CLL (aCLL; n=10) and CLL with Richter transformation (RT-CLL; n=5). We observed an increasing number of CD163+ macrophages expressing FOLR2 between reactive tonsils and CLL categories with increasing aggressivity (Fig. 6A), and demonstrated a significantly higher proportion of FOLR2 expressing CD163+ macrophages in aCLL and RT-CLL compared with cCLL cases (Fig. 6B). Although not statistically significant, the analysis showed a trend toward a higher frequency of FOLR2⁺ NLCs in aCLL and RT-CLL cases (Fig.S6A).

**Figure 6.**
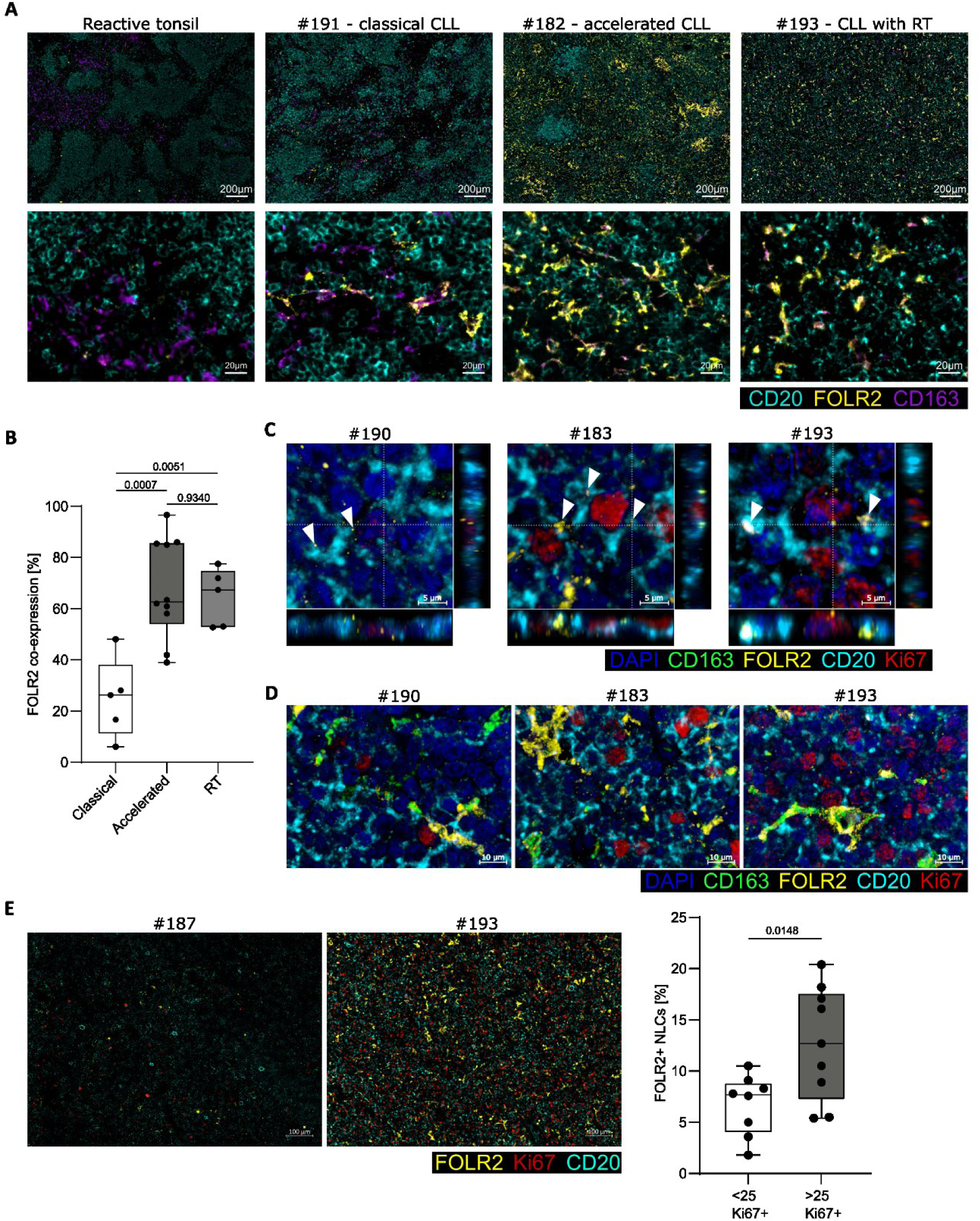
Detection of FOLR2 expression in the lymph nodes of CLL patients. Multiplex immunofluorescence of lymph nodes (LNs) from n = 20 CLL patients (patient characteristic available in Supplementary table S3). Whole FFPE 4µm tissue sections were stained with anti-CD20 (cyan), anti-FOLR2 (yellow), anti-CD163 (purple/green), anti-Ki67 (red; expect #176-178) antibodies, counterstained with DAPI (blue), and visualized with an Axioscan microscope (Fig.6A and E; 20x) or a Zeiss 980 confocal microscope (Fig. 6C and D; Plan-Apochromat, 40x, 1.3 Oil; 3D-projection of 20 z-stacks/0.5µm). **A)** Representative images of a reactive tonsil from healthy donor (n = 1), and patients with: classical CLL (n = 5), accelerated CLL (#183), and CLL with Richter Transformation (#193). Top: large-field images; bottom: magnified regions highlighting a variable level of FOLR2 expression on CD163^+^ macrophages. **B)** Comparison of proportion of FOLR2 expressing CD163^+^ macrophages between CLL groups with increasing aggressiveness: classical CLL (n = 5), accelerated CLL (n = 10), and CLL with Richter Transformation (RT; n = 5), analyzed by ordinary one-way ANOVA. **C)** Detection of FOLR2 signal patches in the plasma membrane (white arrows) or within CD20⁺ cancer cells (patients #190, #183, and #193). Dashed lines indicate regions visualized by full-thickness XZ and XY orthogonal projections. **D)** Detection of FOLR2^+^/CD163^+^ NLCs nearby Ki67^+^ cancer cells. **E)** Left: overview images showing FOLR2^+^ NLCs in relation to Ki67^+^ cancer cells in patients with low (#187) and high (#193) cancer cell proliferation. Right: percentage of FOLR2^+^ NLCs and proliferating Ki67^+^CD20^+^cancer cells were quantified across the whole tissue region. Based on frequency of Ki67^+^ cancer cells, samples were categorized into low (Ki67^+^ CLL cells < 25%; n = 8) and high (Ki67^+^ CLL cells > 25%; n = 9) proliferating groups, using median as a threshold. Statistical comparison of FOLR2^+^ NLCs frequency between the two groups of samples was calculated with unpaired t-test. P-values < 0.05 were considered statistically significant.

Moreover, we detected CD20+ cancer cells containing FOLR2 patches co-localizing with both the plasma membrane, and the cell interior (Fig. 6C, Fig. S6B), indicating the presence of trogocytic cancer cells, previously demonstrated in vitro. Next, we investigated the potential relation between the FOLR2+ NLCs and cancer cell proliferation. We observed that FOLR2⁺ NLCs are closely associated with Ki67⁺ cancer cells (Fig. 6D). Using global image quantification (n = 17), we stratified patient samples into groups with low (<25% Ki67⁺) and high (>25% Ki67⁺) cancer cell proliferation (Fig. 6E), and found that infiltration of FOLR2⁺ NLCs was significantly elevated in the high-proliferation group.

Taken together, these findings confirm the presence of FOLR2⁺ NLCs in patient LNs and demonstrate an association between a higher proportion of FOLR2-expressing macrophages and more aggressive, proliferative cases of CLL. The detection of FOLR2^+^ CLL cells provides evidence of CLL-NLC trogocytosis *in situ*.

## DISCUSSION

Previously, we showed that the M2-like polarization state of NLCs^29^, and their direct interactions with CLL cells, including trogocytosis^17^, are essential in preventing spontaneous apoptosis of cancer cells *in vitro*. Here, we further studied phenotype of NLCs, and their direct interactions with CLL cells, resulting in discovery of FOLR2 as a novel marker of protective NLCs. Moreover, we demonstrated that CLL cells acquire functional form FOLR2 through trogocytosis with NLCs, which is further associated with increased activation and proliferation of cancer cells. Finally, we observed increased proportion of FOLR2 expressing NLCs in the aggressive subtypes of CLL, demonstrated correlation between the percentage of FOLR2**^+^** NLCs and CLL cell proliferation, and detected FOLR2^+^ CLL cells in the patient LNs, collectively supporting the potential relevance of our findings to CLL pathophysiology.

A key finding of our study is the transfer of FOLR2 from NLCs to CLL cells via trogocytosis, revealing a direct mechanistic link between FOLR2⁺ NLCs and enhanced activation and proliferation of cancer cells. FOLR2, thanks to its high affinity, and rapid turnover^25,37,38^, gives CLL trogocytic cells an advantage in folate uptake. Folate is an essential fuel for one-carbon metabolism, impacting nucleic acid, lipids, and amino acids synthesis, necessary for cell growth and proliferation^27^. This would partially explain why FOLR2^+^ trogocytic CLL cells belong to a predominant population of activated and proliferating cancer cells *in vitro,* especially during folate scarcity. Moreover, we demonstrated that the actively cycling population of CLL cells *in vitro* is characterized by a constant level of extracellular FOLR2, which drops as soon as they become quiescent again. This implies that CLL cells engage in trogocytosis with NLCs between each cell cycle and potentially use FOLR2 to sustain their increased folate demands. These findings are supported by our observation that CLL cells remain in close contact with FOLR2^+^ NLCs in the CLL LNs, association between overall infiltration of FOLR2^+^ NLCs and more proliferative disease, and detection of FOLR2^+^ cancer cells *in situ*. Moreover, it was reported and that during CLL progression, NLCs show an increased infiltration into proliferation centers^4^, hence being more accessible for the actively proliferating CLL cells. In the literature, there are similar examples of trogocytic cells, gaining increased adaptability. For instance, Lu et al. demonstrated that NK cells can acquire a tyrosine kinase receptor TYRO3 from tumor cells, resulting in their increased activation and proliferation^39^.

Moreover, our data indicate an unexplored bias in protein transfer during trogocytic CLL-NLC interaction. Out of the tested proteins, FOLR2 is the main NLC-derived protein detectable on CLL cells in the resting conditions. FOLR2 is a small, GPI-anchored protein, located on the plasma membrane. In contract, CD16, CD163, and CD206, are all integral transmembrane proteins with stronger membrane association^48^, suggesting that observed trogocytosis primarily involves the outer leaflet of NLC plasma membrane. Interestingly, CD14, also a small GPI-anchored protein, predominantly co-expressed with FOLR2, showed much smaller transfer, and retention on CLL cells, which further indicates the selectivity and significance of FOLR2 transfer. Moreover, upon CD40L+IL-15 priming, we observed not only a 10-fold increase in the transfer of FOLR2 to cancer cells, but also a certain acquisition of CD11b and CD14, indicating that activation is intensifying interaction between CLL cells and NLCs, and thus the trogocytosis process itself. Given that FOLR2 reportedly inhibits CD11b/CD18-mediated macrophage adhesion^49^, and decreased adhesion could promote trogocytosis in macrophages^50^, we suggest that FOLR2 modulates CLL–NLC contact dynamics, also explaining its preferential acquisition by CLL cells.

Importantly, with this study, we defined FOLR2 as the ultimate marker of NLCs, allowing us to discriminate between protective and non-protective phenotypes *in vitro.* Moreover, we confirmed FOLR2 expression in CLL patient LNs, and observed an increased proportion of FOLR2⁺CD163⁺ NLCs in more progressive CLL cases. These results highlight previously unknown heterogeneity of NLCs, stressing the importance of further characterization of these peculiar cells.

While we demonstrated a clear association between FOLR2 expression and protective phenotype of NLCs, the exact role of FOLR2 in functionality of these cells remains speculative. In macrophages, FOLR2 expression has been proposed to promote folate accumulation, and stabilization of immunosuppressive phenotype through epigenetic changes^28^. This aligns with findings that NLCs are profoundly immunosuppressive and promote T-helper cell expansion, thereby weakening antitumor immunity ^24^. We further speculate that FOLR2^high NLCs might act as folate reservoirs for proliferating CLL cells, particularly during phases of rapid disease expansion when systemic folate availability becomes limiting ^28^. Although FOLR2 acquisition partially explains the phenotype of trogocytic CLL cells, additional molecules transferred from NLCs likely contribute to enhanced activation and proliferation. Trogocytosis could support CLL cell activation through acquisition of activating receptors or facilitating cross-presentation of autoantigens^46^ from NLCs. Moreover, trogocytosis with NLCs could help CLL cells to optimize their metabolic requirements through acquisition of lipids and other biomolecules required for cell proliferation^20,47^. Moreover, we do not exclude impact of other mechanisms to be involved in beneficial effects of NLCs, including release of soluble factors or extracellular vesicles (EVs). Nevertheless, our previous findings showed that transwell cultures, permissive to soluble factors and EVs, abolish most supportive impact of NCLs^19^. Moreover, we observed a very strong association of CLL cells with NLCs following CD40L + IL-15 stimulation (Fig. S4E). Finally, if EVs were contributing to FOLR2 acquisition by CLL cells, the dynamic fluctuation of FOLR2 levels across cell-cycle would be unlikely.

This work underlies the central role of FOLR2^+^ NLCs in supporting CLL cells, and characterizes a novel, trogocytosis-mediated type of pro-tumoral interaction between these two cell types. We believe that uncovering the intricacy of trogocytosis in CLL could increase our understanding of the cancer cell interactions within the TME, leading to more efficient therapeutic strategies, potentially relevant for other types of cancers.

## Acknowledgments

The authors thank all the patients who generously donated clinical samples for this study. This work was supported in part by the Janssen Foundation, Région Midi-Pyrénées, La Ligue Nationale Contre le Cancer, CNRS, and Toulouse III University. We would like to thank the members of the platform IMAG’IN (l’Institut Universitaire du Cancer de Toulouse), especially N. Van Acker, FX. Frenois and A. Belin-Artola for the IHC staining of LN. We are grateful to P. Gravelle for facilitating communication with the Anatomo-Pathology Department of the IUCT Oncopole and to A. Quillet-Mary for her critical review of the manuscript. Schematics were created using BioRender.com.

## Institutional Review Board Statement

The study was conducted according to the guidelines of the Declaration of Helsinki, and approved by the Institutional Review Board (or Ethics Committee) of INSERM (protocol code DC-2013-1903 and date of approval 2013).

## Authorship

Contribution: M.D. designed the study, performed the experiments, interpreted data, and wrote the manuscript; C.B. processed the healthy-donor blood samples; B.G. participated in design of the research; L.Y. provided patients’ blood samples and associated clinical data; C.L. provided the patients’ lymph node samples, associated clinical data, and supervised IHC staining; V.P. provided the funding; M.P provided the funding, designed the study, interpreted data, and wrote the manuscript. All the authors critically reviewed the manuscript.

