## Supplementary material for "Trogocytosis-mediated transfer of FOLR2 from nurse-like cells to CLL cells is associated with their activation and proliferation": Supplement_091225_final.docx

**Supplementary methods:**

### Cell isolation

Peripheral Blood Mononuclear Cells (PBMC) were isolated from blood of CLL patients or HD buffy coats by density-gradient centrifugation, and used directly for the downstream processing or cryopreserved using CryoStor CS10 (STEMCELL).

CLL cells were purified from PBMC, using a EasySep™ Human B Cell Enrichment Kit II Without CD43 Depletion negative selection kit (STEMcell, France), according to manufacturer's protocol. CLL cells used for heterologous co-cultures were purified from cryopreserved PBMC. To minimize cell death, PBMC were thawed the day before purification, and cultured in appropriate media at 10 × 10^6^ c/mL. For experiments measuring cancer cell proliferation, right after purification, CLL cells were stained with CellTrace™ Violet (CTV; Invitrogen) according to manufacturer instructions. Cell purity and viability were evaluated by flow cytometry using CD5 and CD19 antibodies and viability staining, and consistently exceeded 95% across all samples (data not shown).

Healthy donor monocytes were purified using CD14 MicroBeads positive selection kit and MS/LS columns (Miltenyi Biotec, France), according to the manufacturer’s protocol. Isolation efficacy was evaluated by flow cytometric analysis of CD14⁺ cell frequency, and exceeded 95% (data not shown). Freshly isolated monocytes were used to generate monocyte-derived macrophages (MDMs) or healthy donor NLCs (HD-NLCs).

### Cell cultures

*In vitro* cultures were conducted in complete medium, composed of RPMI 1640 with GlutaMAX (Gibco, France), supplemented with 10% fetal bovine serum (FBS, Life Technologies, France) and 100 µg/ml of penicillin/streptomycin (Sigma-Aldrich, France). For experiments with varying concentration of folic acid (FA), media were prepared on the basis of RPMI 1640 without folic acid (Gibco, France) and 100 µg/ml of penicillin/streptomycin. For FA^high^ condition, the media was supplemented with standard 2.27μM FA concentration. Subsequently, both FA^high^ and FA^low^ media were supplemented with 10% FBS. For FA^negative^ condition, 10% of dialyzed FBS (Sigma, Germany) was used instead.

Cells were plated in tissue culture-treated plates (Corning, USA) at varying volumes: 4 mL per well for 6-well plates, 1.5 mL per well for 12-well plates, and 500 µl per well for 48-well plates. All cultures were maintained at 37°C in a humidified atmosphere with 5% CO2.

Autologous nurse-like cells (NLCs) were generated by the culture of freshly isolated CLL PBMC at 10 × 10^6^ cells/mL in complete medium for up to 15 days^1^. For NLCs phenotyping, combined fractions of floating cells collected by pipetting, and adherent cells detached by Accutase were used.

Monocyte-derived macrophages (MDMs) were generated by activating purified CD14⁺ monocytes with either 20 ng/mL CSF-1 or 50 ng/mL GM-CSF during 6 days for initial differentiation of M2-like or M1-like MDMs, respectively. Subsequently, cells were polarized for 48 hours with 25 ng/mL IL-4 and 25 ng/mL IL-13 to generate M2-like MDMs, or with 100 U/mL IFNγ and 10 ng/mL LPS to generate M1-like MDMs.

**Heterologous co-culture system**

A heterologous co-culture system combining purified monocytes from healthy donors (HD) and CLL cells from patient blood was used for generation of HD-NLCs, and studying FOLR2 trogocytosis in conditions mimicking lymph node microenvironment.

Basic protocol (fig. S4A): CD14⁺ healthy donor monocytes were plated at 0.5 × 10⁶ cells/mL in a 48-well plate and cultured in 500 μL of complete medium (CM) supplemented with 20 ng/mL CSF1. After 6 days, co-culture was initiated by removing the spent medium and adding purified naïve CLL cells at 10 × 10⁶ cells/mL in 500 μL of fresh CM. After 5 days of co-culture, cells were activated with either 10 ng/mL IL-2 and 1 μg/mL CpG, or 50 ng/mL CD40L and 10 ng/mL IL-15, for 72 hours.

Optimized protocol (fig. 4A): MDMs were generated by plating CD14⁺ cells at 1 × 10⁶ cells/mL in 250 μL of CM supplemented with 20 ng/mL CSF-1, in a 48-well plate. After 6 days, medium was replaced and purified naïve CLL cells were added at 6 × 10⁶ cells/mL in 250 μL of CM, to initiate the co-cultures. 4 days later, after polarization of HD-NLC, 250 μL of fresh medium containing CD40L was added to a final concentration of 25 ng/mL. 2 days later, 300 μL of medium was replaced, and cells were further stimulated with 50 ng/mL CD40L and 10 ng/mL IL-15 for 5 days.

### Flow cytometry

### Extracellular staining: All tested samples were re-suspended in the staining buffer (5% FBS, 0,5% BSA, 2mM EDTA, 0.1% sodium azide in PBS), supplemented with 2.5 µg/mL Human BD Fc Block™ (BD Pharmingen, France) and 1% human serum to prevent the non-specific antibody binding, and incubated for 15 min at 4°C. Subsequently, cells were stained with antibodies at saturating concentrations, for 20 min at 4°C. After washing, cells were re-suspended in PBS, containing 0.8 µg/mL 7-AAD (Sony, USA).

Intracellular staining: Cells were stained with FVS620 viability dye (BD) according to manufacturer’s protocol. Following the extracellular staining, cells were fixed and permeabilized with True-Nuclear™ Transcription Factor Buffer Set (Biolegened). Next, cells were stained with anti-Ki67 mAb (Invitrogen) for 20 min at RT, washed and re-suspended in PBS.

AnnexinV/7-AAD staining of CLL cells: Cells were washed and resuspended in Annexin-V buffer and stained with Annexin-V-FITC (Miltenyi, Germany) according to manufacturer protocol. Afterward, cells were resuspended in Annexin-V buffer containing 0.8 µg/mL 7-AAD.

Results: were analyzed using FlowLogic 7.00.2 software (Inivai Technologies). Gating strategies for the analysis of CLL cell activation and proliferation are provided in the Supplementary Information (Fig. S4E and S5A, respectively). Gates for FOLR2⁺ cells were set using fluorescence-minus-one (FMO) controls.

The analysis of the enrichment of FOLR2^+^ trogocytic cells in the population of activated CLL cells, was made by introducing a FOLR2 enrichment score (ES_FOLR2_), further correlated with the total percentage of CLL cells positive for a specific activation marker (Fig. S4F; A). Ki67 enrichment score was calculated analogously, to represent enrichment of Ki67^+^ CLL cells in the population of FOLR2^+^ trogocytic CLL cells.

**Gene expression analysis with quantitative RT-PCR**

To evaluate FOLR2 gene expression, cell pellets stored in -80°C, were resuspended in TRIzol™ Reagent, and RNA was isolated using Direct-zolTM RNA MiniPrep kit according to manufacturer instruction. Subsequently, reverse transcription was performed on purified RNA using SuperScript™ III enzyme, and complement DNA was amplified by polymerase chain reaction (PCR). Primers targeting FOLR2 gene were based on^2^, and validated using M2-like MDMs and FOLR2 KO M2-like MDMs as positive and negative controls respectively, prior to application in CLL samples. Quantitative PCR using PowerUp™ SYBR™ Green Master Mix, was performed on StepOnePlus Real-Time PCR Systems (Applied Biosystem), using standard protocol with Tm = 60°C and 40 amplification cycles. FOLR2 gene expression was calculated using the comparative 2⁻ΔΔCt method, normalizing the results with GADPH housekeeping gene.

### FOLR2 gene knockout using RNP Crispr-Cas9 method

Freshly isolated CD14^+^ monocytes from a healthy donor were activated with 20 ng/mL CSF-1, and seeded in CM without antibiotic for 16h. CrRNA and tracrRNA were reconstituted in Nuclease Free Duplex Buffer (IDT, USA), and mixed in equimolar concentrations, followed by heating at 95°C for 5 min. Each gRNA was combined with Alt-R™ S.p. HiFi Cas9 Nuclease V3 at a 1:1.2 molar ratio, incubated at 37°C for 15-20 min, and cooled down to RT. After detachment, monocytes were resuspended in buffer T (Neon™ Transfection System 100 μL Kit), mixed with RNP complexes, and electroporated in Neon Tip with Neon Transfection System, at 1900V, 20ms, single pulse. Afterwards, cells were plated in antibiotic free medium with 25 ng/mL CSF-1, and cultured for 5 days to allow maturation of MDMs. After confirmation of FOLR2 gene knockout (KO) effectiveness by qPCR (Fig. S2B) or flow cytometry (Fig. S2D), naïve purified CLL cells were mixed with FOLR2 KO MDMs to induce polarization of HD-NLCs and facilitate cell-cell interactions. After 6 days, CLL cells were analyzed by flow cytometry.

**Immunofluorescence of *in vitro* cells**

Following the long-term *in vitro* culture (12-14 days), CLL PBMC were harvested and re-suspended in the staining buffer (5% FBS + 0.5% BSA + 1mM EDTA in PBS) containing 1% human serum and 2.5 µg/mL Human BD Fc Block™ (BD Biosciences, France) and incubated for 15 min at 4°C. For detection of extracellular antigens, cells were first stained indirectly with rabbit anti-FOLR2 mAbs (Abcam) for 20 min at 4°C, followed by washing wish PBS, and incubation with goat anti-rabbit IgG secondary antibodies conjugated with AF647 (Invitrogen) for 15 min at 4°C. Next, cells were incubated with mouse anti-CD19 antibody conjugated with BV421 (Sony) for 20 min at 4°C. Subsequently, cells were washed four times with PBS and fixed with 4% PFA in PBS for 20 min at RT. After three washes with PBS, cells were counterstained with DAPI (1μg/mL) for 5 min, washed twice with PBS and left on the cover glass O/N at 4°C to facilitate attachment of the cells. The next day, slides were mounted using Fluoromount media (Thermo Fisher Scientific), and imaged using confocal microscope LSM 780 or 880 (Zeiss). Images were analyzed using ZEN blue 3.12 software (Zeiss).

**Immunohistochemistry of Patient Tissue Samples**

Multiplex Immunofluorescence (mIF): the BOND RX staining platform (LEICA Biosystems, Germany) was used to automate the mIF staining procedure on 4 µm FFPE tissue sections. After baking and dewaxing, tissue slides were heat-pretreated for 20 min at 100°C using ER2 pretreatment solution (pH9, LEICA Biosystems, Germany). The slides were blocked for endogenous peroxidase activity using the Discovery inhibitor (15 minutes at room temperature) (07017944001, Roche Diagnostics, Switzerland). The slides were then stained for 5-plex immunofluorescence using the OPAL^TM^ Technology (AKOYA Biosciences, USA) with sequential denaturation using ER1 pretreatment buffer (pH6, LEICA Biosystems, Germany), for 20 min at 97°C between each antibody-staining cycle. Primary antibodies were applied and incubated during 30 min at room temperature programmed in the aforementioned staining cycles in the following order: FOLR2 (clone EPR25731-70 (ABCAM Netherlands), at final dilution of 1/200, Envision Flex antibody diluent, Agilent Technologies), Ki67 (clone SP6, Ready to use, Zytomed, by Diagomics, France), CD163 (clone MRQ-26, Ready to use, Roche Diagnostics), CD68 (clone PGM-1, at dilution of 1/100, in Envision Flex antibody diluent) and CD20 (clone L26, Ready to use antibody diluted at ¼ in Ventana Antibody diluent, Roche Diagnostics). Subsequently, slides were incubated using the Opal polymer HRP detection system, allowing for OPAL^TM^ dye precipitation. Five OPAL^TM^ dyes with excitation and emission wavelengths compatible with our whole slide imaging system were used: OPAL^TM^ Polaris 480, OPAL^TM^ 520, OPAL^TM^ 570, OPAL^TM^ 620 (1/300 in 1x Plus Automation Amplification Diluent, AKOYA Biosciences, USA) and OPAL^TM^ 690 (1/150 in 1x Plus Automation Amplification Diluent). The tissue slides were counterstained using Spectral DAPI (AKOYA Biosciences, USA) and mounted with Invitrogen^TM^ ProLong^TM^ Gold Antifade Mounting medium (Life Technologies, USA). Whole tissue slides were scanned with ZEISS Axio Scan.Z1 microscope, using a Plan-Apochromat 20x/0.8 M27 objective in 1,1 binning mode. For visualization of trogocytosis in the tissue, selected regions were imaged with Zeiss 980 confocal microscope, using Plan-Apochromat 40×/1.3 oil objective.

Image analysis: basic image processing and analysis, including region-of-interest selection and co-localization measurements, were performed using ZEN 3.12 software (ZEISS, Germany). For signal quantification, the mIF images were initially processed with HALO software (version 3.6, Indica Labs, USA). Briefly, each section was manually annotated to include the maximal tissue area while excluding artifacts introduced during sample preparation. Subsequently, segmentation parameters were optimized to achieve the most consistent cell detection, and threshold for each marker was adjusted to minimize false-positive events. The annotated regions were then segmented, and the resulting image-analysis data were exported as FCS files for downstream analysis in Flow Logic 700.2A software (Inivai Technologies, Australia). Each cell was classified based on the combination of specific dye intensities and the completeness of its nuclear or cytoplasmic region. Results were analyzed and plotted using GraphPad Prism 10.1.2 (USA).

**Supplementary Figures**

**Supplementary Figure S1**

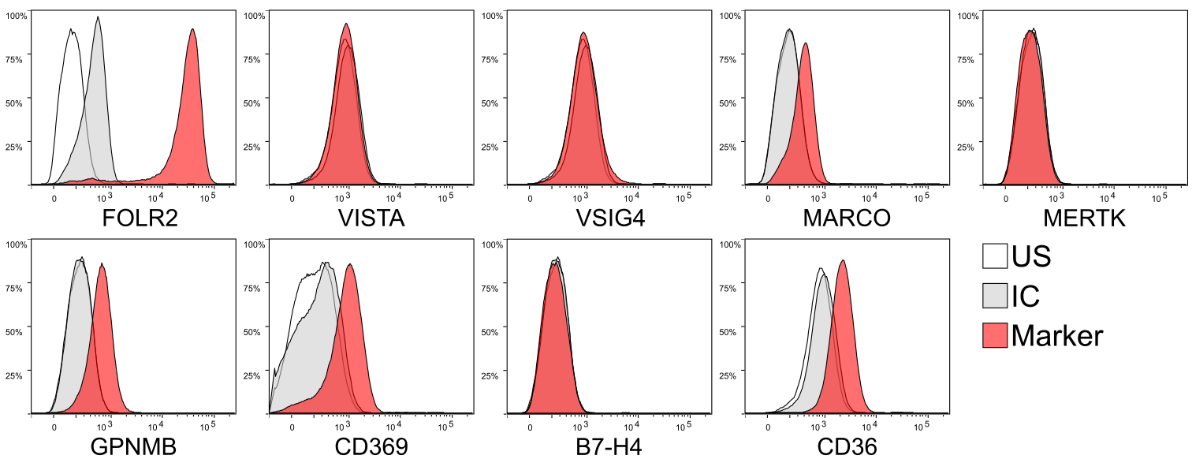

**Figure S1A. Screening of novel potential marker for NLCs.** NLCs were generated by high-density culture of CLL PBMC. After 14 days, cells were collected, stained with specific antibodies, and analyzed by flow cytometry. NLCs were gated based on high forward (FSC) and side scatter channel (SSC), and results for each marker were depicted by representative histogram overlay between signal from: unstained cells (US), isotype control antibody (IC), and antibody against antigen of interest (Marker).

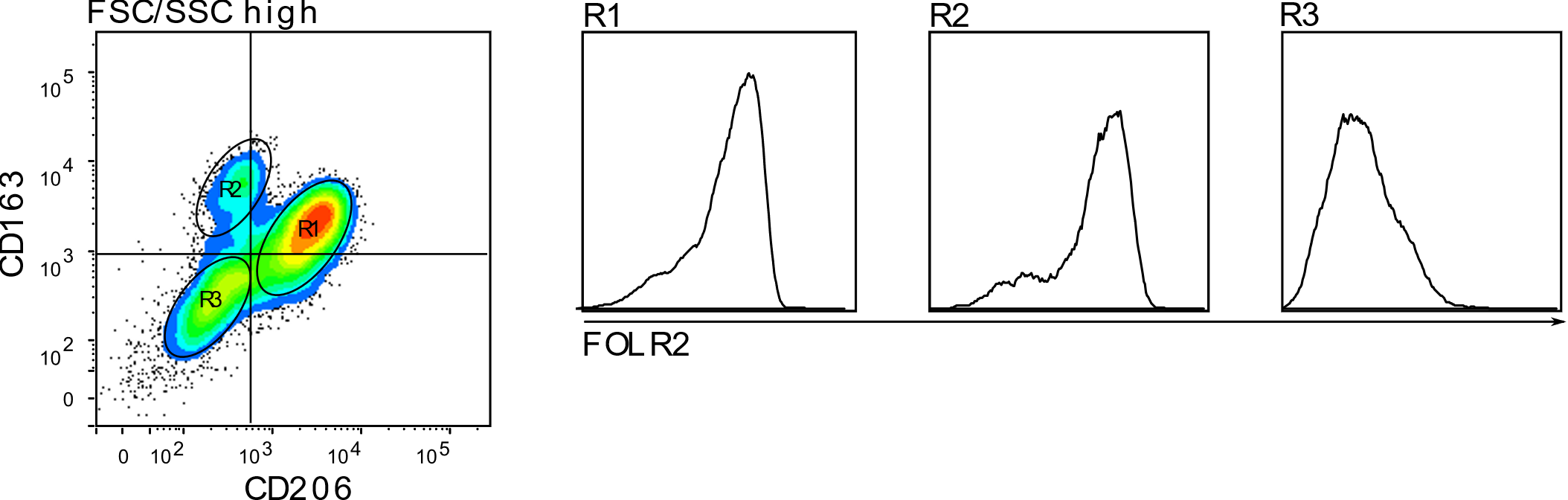

**Figure S1B. Detection of heterogeneous subpopulations of NLCs with varying expression of FOLR2.** NLCs generated *in vitro* were analyzed by flow cytometry. An exemplary dot plot shows NLCs, divided into three subpopulations (R1-R3), based on the expression of typical macrophage’s markers: CD163 and CD206.

| FOLR2^high^ group | FOLR2^neg/low^ group |
| --- | --- |

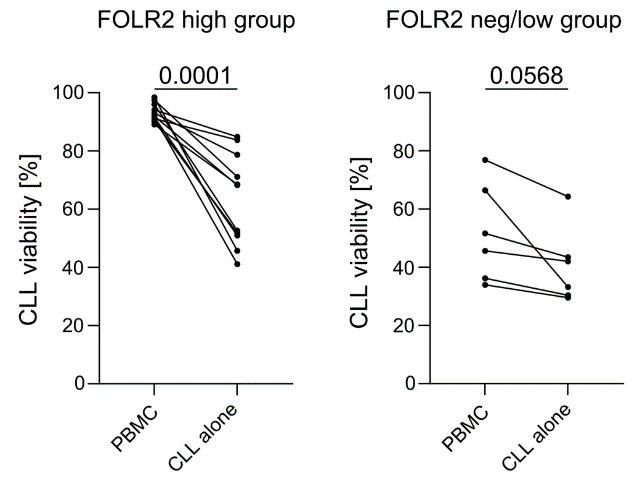

**Figure S1C. Presence of FOLR2^high^ NLCs is associated with increased survival of CLL cells *in vitro*.** For a subset of the patient samples described in Fig. 1F-G, in addition to PBMC cultures (which allow for the outgrowth of NLCs), a fraction of cancer cells was purified by negative selection and cultured at the corresponding density of 10 × 10⁶ cells/mL (CLL alone). CLL cell viability was assessed by Annexin V/7-AAD staining between days 12–14 for PBMC cultures and on day 7 for CLL cultured alone. Paired PBCM-CLL alone samples from the same patient were divided into two groups (FOLR2^high^ with n = 11 samples, and FOLR2^neg/low^ with n = 6 samples), based on FOLR2 expression by NLCs (according to Fig. 1F). Survival of CLL cells between the two culture conditions was compared using a paired, two-tailed t-test, with p-values < 0.05 considered statistically significant.

**Supplementary Figure S2**

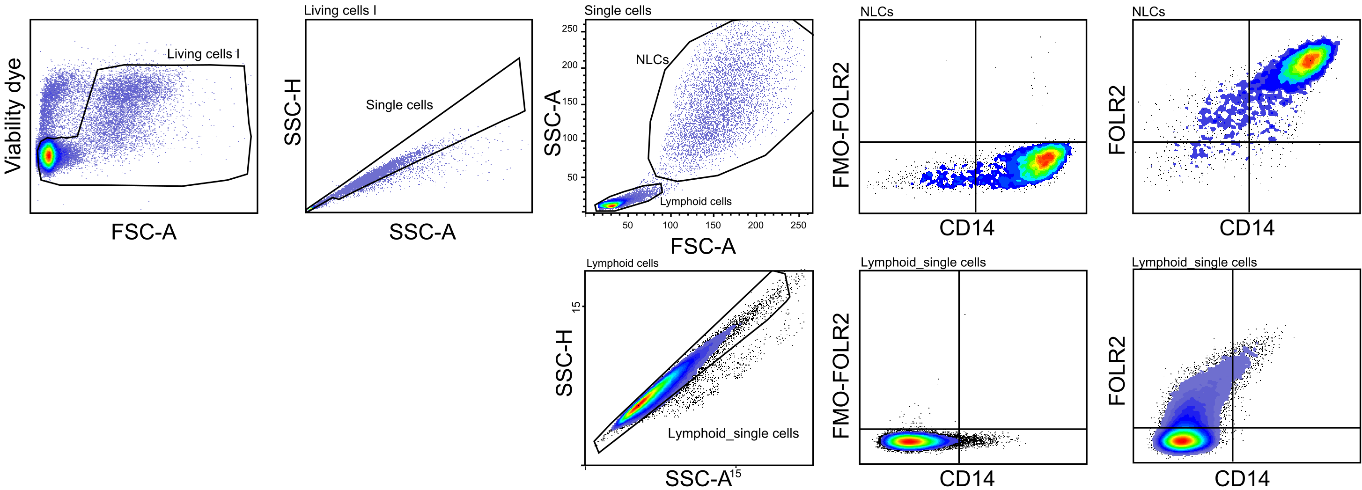

**Figure S2A. Gating strategy for FOLR2 signal measurements on NLCs and lymphoid cells.** Dead cells were identified based on 7-AAD staining, gating was adjusted to account for high autofluorescence of NLCs. Cell doublets were excluded based on forward (FSC, not shown) and side-scatter (SSC). PBMCs were divided into NLCs and lymphoid cells based on FSC and SSC difference. Gating of the FOLR2 signal was determined using the corresponding Fluorescence-minus-one (FMO) control.

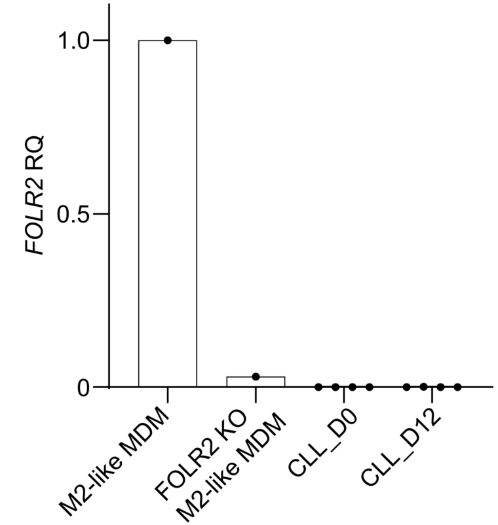

**Figure S2B. CLL cells do not express the FOLR2 gene.** Freshly isolated CLL PBMCs were divided into two parts: 1) used to purify CLL cells by negative selection, followed by freezing the cancer cells in −80°C (CLL_D0); 2) was cultured *in vitro* for 12 days to allow NLC outgrowth, after which CLL cells were purified by negative selection and frozen at −80°C (CLL_D12). After collecting all samples, FOLR2 gene expression in CLL cells was measured by qPCR, using M2-like MDMs and *FOLR2* KO M2-like MDMs as positive and negative control respectively. The graph displays pooled data from n = 4 independent experiments and four different patient samples, and M2-like MDM controls from a single healthy donor. Results of the analysis were represented as Relative Quantification (RQ) according to FOLR2 expression by M2-like MDMs.

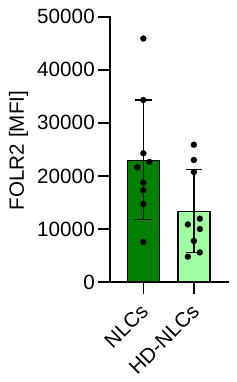

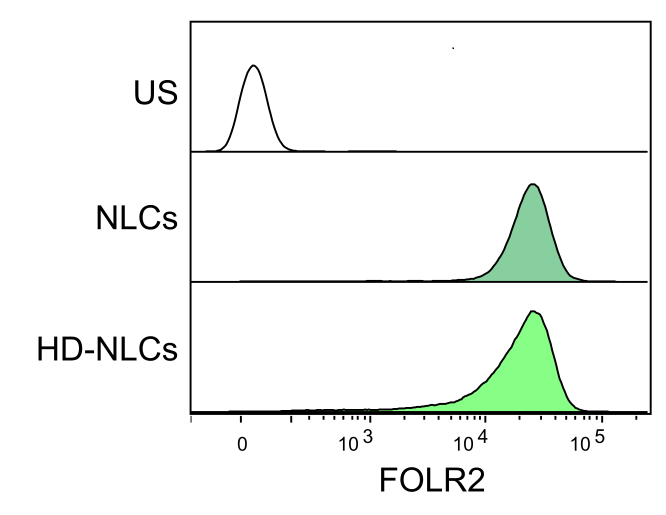

**Figure S2C. HD-NLCs express comparable levels of FOLR2 to autologous NLCs.** Flow cytometry analysis of FOLR2 expression on NLCs (dark green) generated through standard autologous CLL PBMC culture, and healthy-donor NLCs (HD-NLCs; light green), obtained through heterologous co-culture of HD CD14^+^ monocytes and purified CLL cells. US – unstained NLCs control. Left – histogram overlay of a representative experiment; right – bar plot displaying results from n=9 different patients (NLCs) and healthy donors (HD-NLCs). Bars represent mean ± SD. HD-NLCs have a tendency to express slightly lower levels of FOLR2 as compared with NLCs, nevertheless, no statistical significance was detected using unpaired t-test.

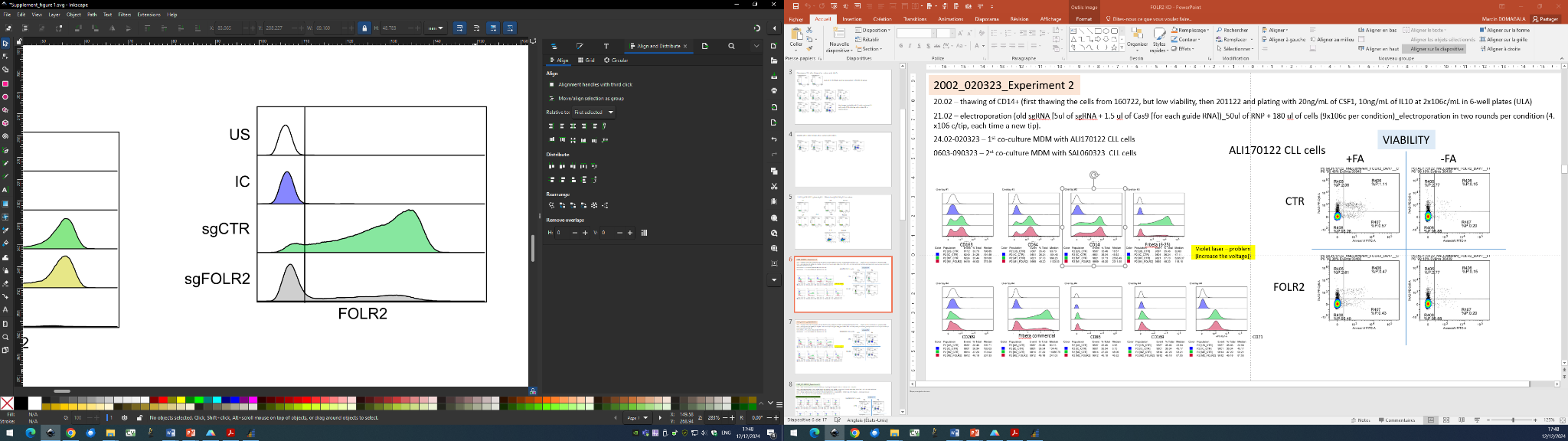

**Figure S2D. Downregulation of FOLR2 protein in MDMs following *FOLR2* knock-out.** CD14^+^ monocytes from healthy-donor PBMC were isolated by positive selection, and activated with CSF1 for 16h. Subsequently, the monocytes were electroporated with sgRNA targeting FOLR2 with RNP Crispr-Cas9 approach. On day 5 after electroporation, monocyte-derived macrophages (MDMs) were collected, and FOLR2 expression was measured by flow cytometry. Results are represented as histogram overlay. US - unstained cells, IC - isotype control, sgCTR - MDMs after electroporation with a control sgRNA, sgFOLR2 – MDMs after electroporation with sgRNA targeting *FOLR2*.

**Supplementary Figure S3**

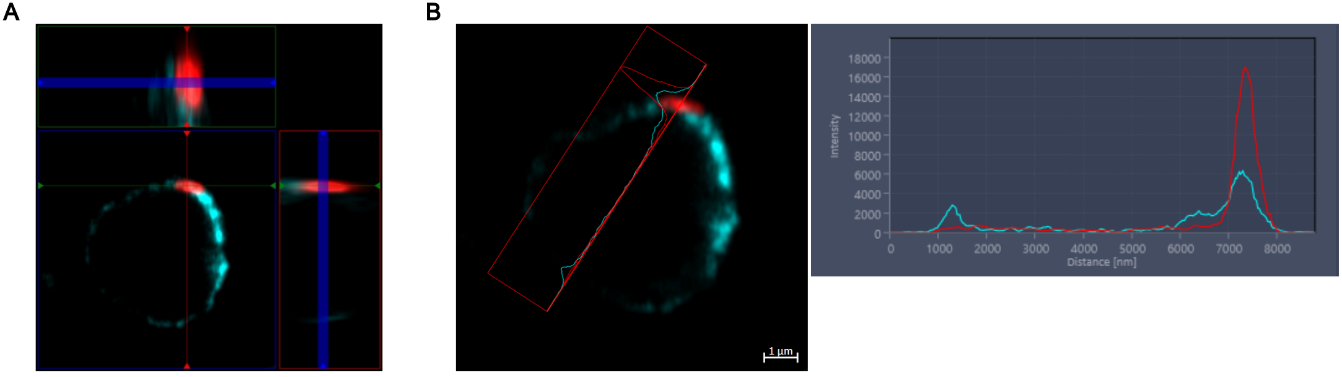

**Figure S3A. FOLR2 co-localizes with CD19 in the plasma membrane of CLL cells.** Visualization of signal co-localization between CD19 (cyan) and FOLR2 (red) in the plasma membrane of CLL cells after co-culture with NLCs. Results are presented as orthogonal projection (A) and co-localization profile (B).

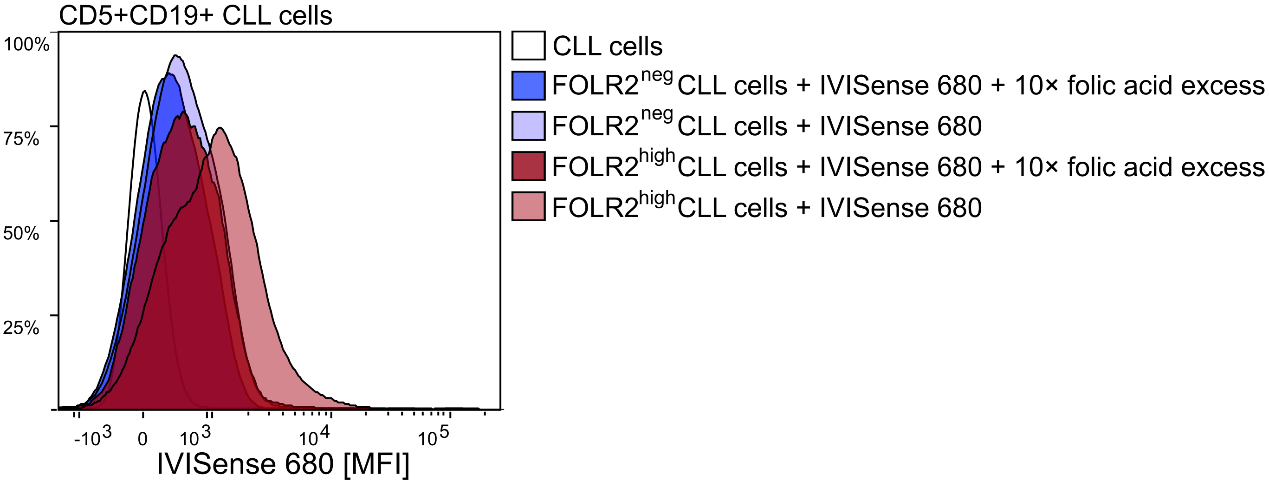

**Figure S3B. FOLR2^+^ trogocytic CLL cells exhibit increased folic acid acquisition.** Flow cytometry analysis of fluorescent folic acid probe (IVISense 680) acquisition by CLL cells. CLL cells following co-culture with NLCs, were re-plated and re-suspended in HBSS buffer supplemented with 5% of dFBS, and incubated with IVISense680 for 90 min, cell culture incubator at 37°C. IVISense 680 signal was analyzed by flow cytometry, following gating on FOLR2^neg^ (blue) or FOLR2^high^ (red) subpopulation of CLL cells. As a control of IVISense680 specificity, cells were incubated with 10x excess of unlabeled folic acid. Representative results from n = 2 independent experiments.

**Supplementary Figure S4**

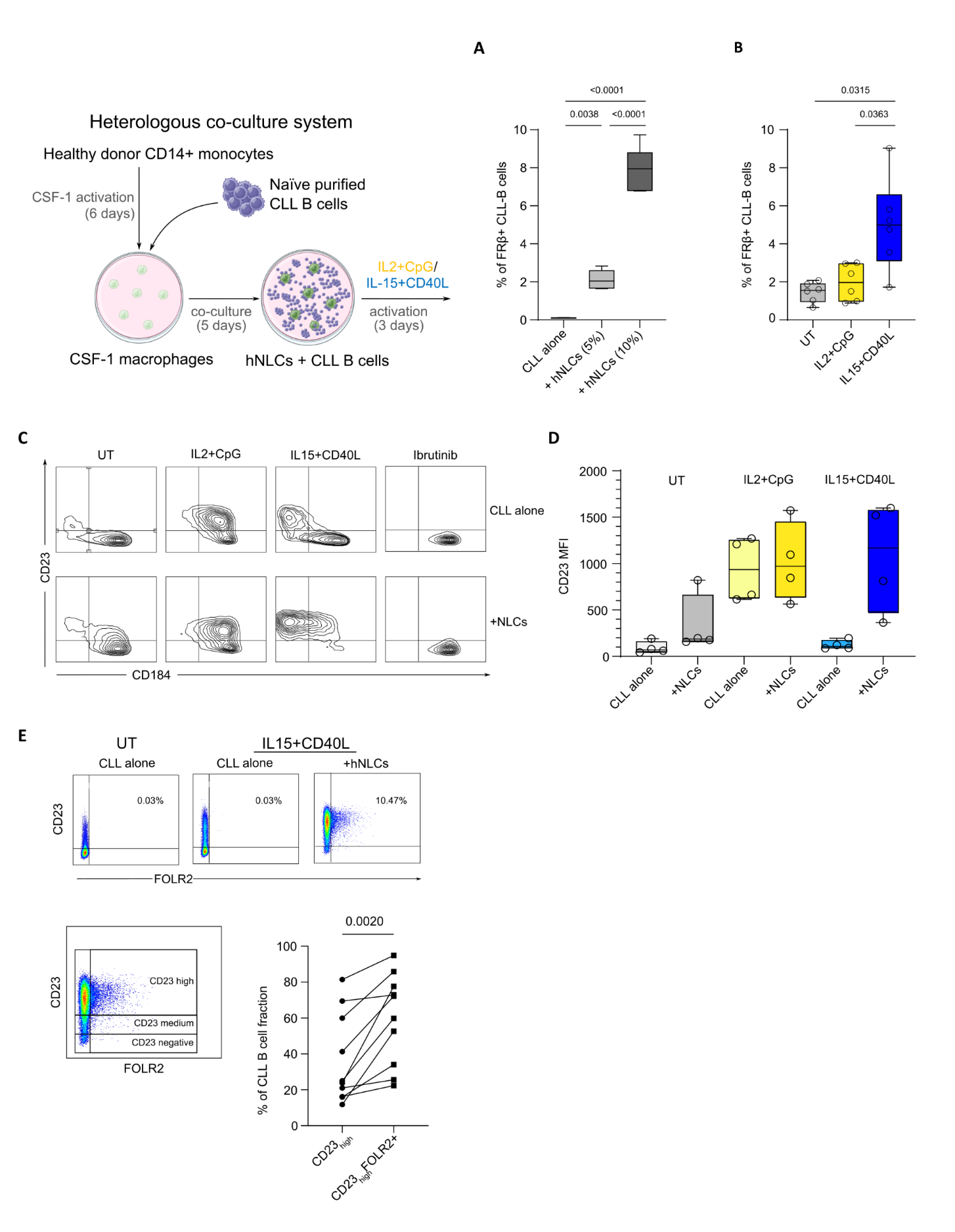

**Figure S4A.** Schematic of the heterologous co-culture system used to compare the effects of CD40L+IL-15 and IL-2+CpG activation protocols.

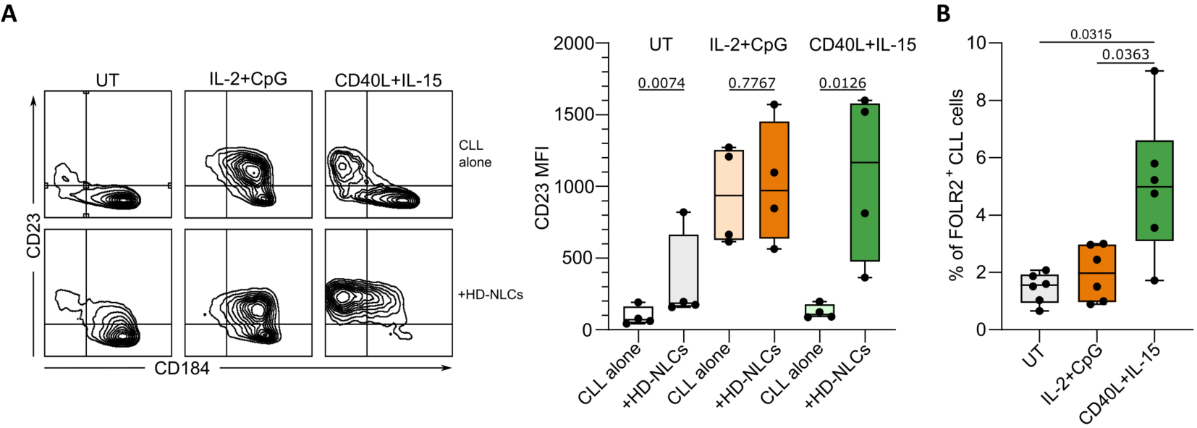

**Figure S4B.** **CD40L+IL-15 leads to potent, NLC-dependent CLL cells activation and increased FOLR2 acquisition.** CLL cells were cultured alone or in a heterologous co-culture system, followed by priming with IL-2+CpG or CD40L+IL-15 for 3 days. CD23, CD184 and FOLR2 signals on CLL cells were measured by flow cytometry. A) Representative density dot plots of CD23 and CD184 expression on CLL cells depending on activation protocol and presence of HD-NLCs; ratio paired t-test comparison of CD23 expression between CLL cultured alone or with HD-NLCs for the listed activation conditions, based on data from n = 4 different patient samples. B) Impact of IL-2+CpG or CD40L+IL-15 treatment on FOLR2 acquisition by CLL cells in heterologous co-cultures. Data from n = 6 different patient samples was analyzed using repeated measures ANOVA with Geisser-Greenhouse correction, with P-values < 0.05 considered statistically significant.

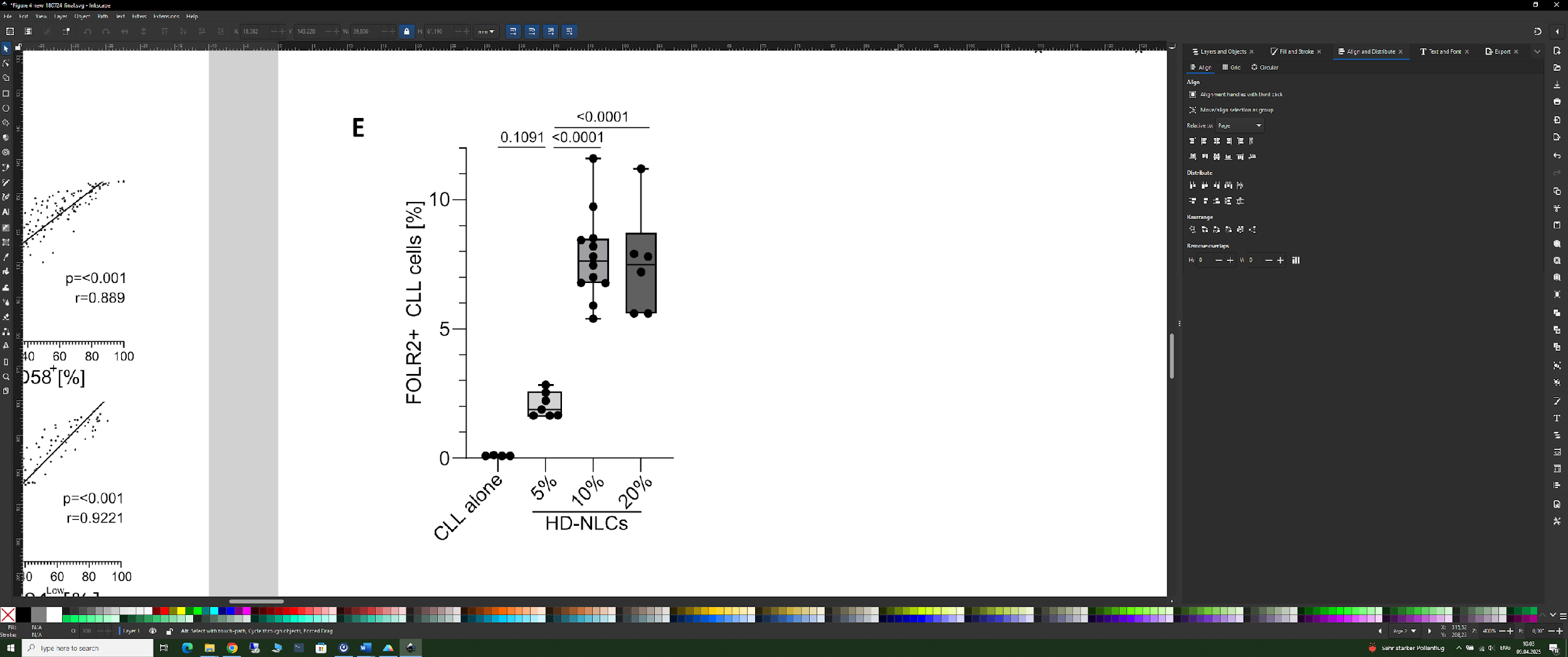

**Figure S4C. Effect of HD-NLC proportion on FORL2 acquisition by CLL cells.** Monocytes were plated with increasing percentage in relation to CLL cell number used for the subsequent co-cultures. After 6 days of CSF1 activation, the media was replaced, and co-cultures were initiated by addition of freshly purified CLL cells at 6 × 10^6^c/mL. After 5 days of co-culture, the level of FOLR2 on CLL cells was measured by flow cytometry. Results from n = 2 independent experiments (samples from 4 (CLL alone) or 6 different patients) were analyzed with ordinary one-way ANOVA, with p-values < 0.05 considered statistically significant.

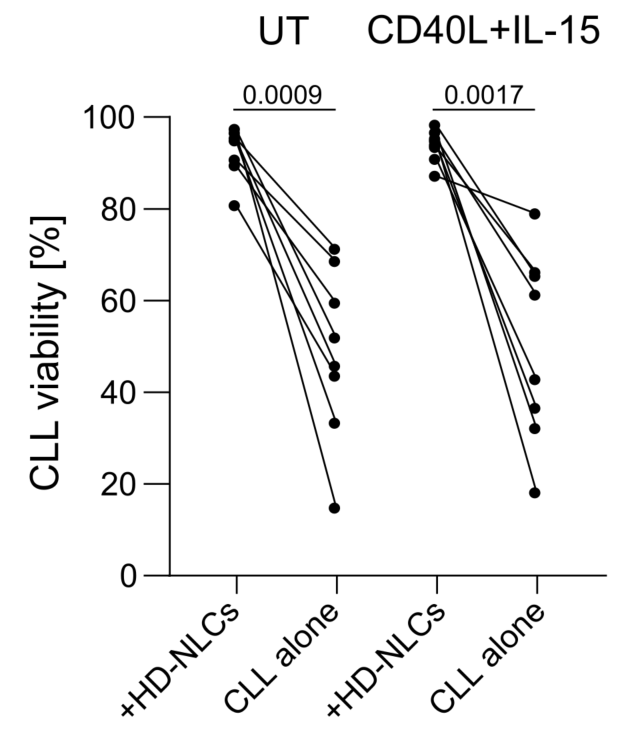

**Figure S4D. HD-NLCs protect CLL cells from spontaneous apoptosis in vitro, independent of CD40L+IL‑15 activation.** CLL cells from heterologous co-cultures (+HD-NLCs) or monocultures (CLL alone) were treated with CD40L + IL-15 according to the optimized activation protocol. After 5 days, CLL cell viability was assessed by Annexin V/7-AAD staining (n = 1 independent experiment; samples from 8 different patients). CLL cell survival between co-cultures and monocultures, within each experimental condition were analyzed using a paired, two-tailed t-test, with p-values < 0.05 considered statistically significant. UT – untreated cells.

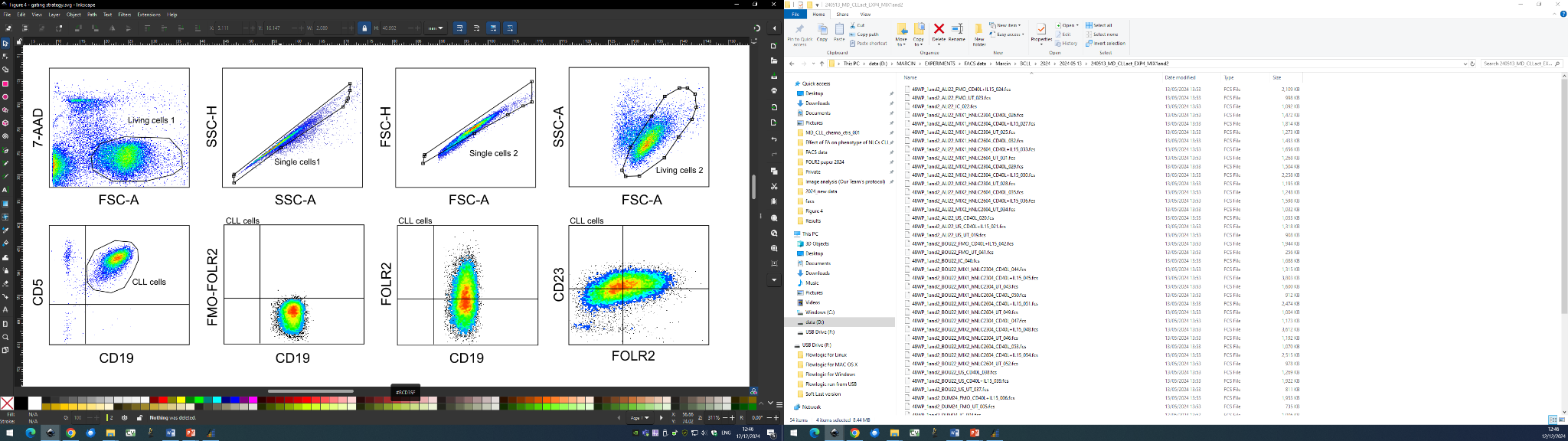

**Figure S4E. Gating strategy for evaluation of CLL cell activation and FOLR2 acquisition.** Dead cells were identified based on positive staining with 7-Aminoactinomycin D (7-AAD) and forward scatter (FSC). Cell doublets were excluded based on FSC and side-scatter (SSC) parameters. Remaining apoptotic/dead cells that showed not signal from 7-AAD were eliminated based on combination of FSC and SSC signals. CLL cells were gated based on positive staining with anti-CD5 and CD-19 antibodies. Positive gate for the FOLR2 signal was determined using the corresponding Fluorescence-minus-one (FMO) control.

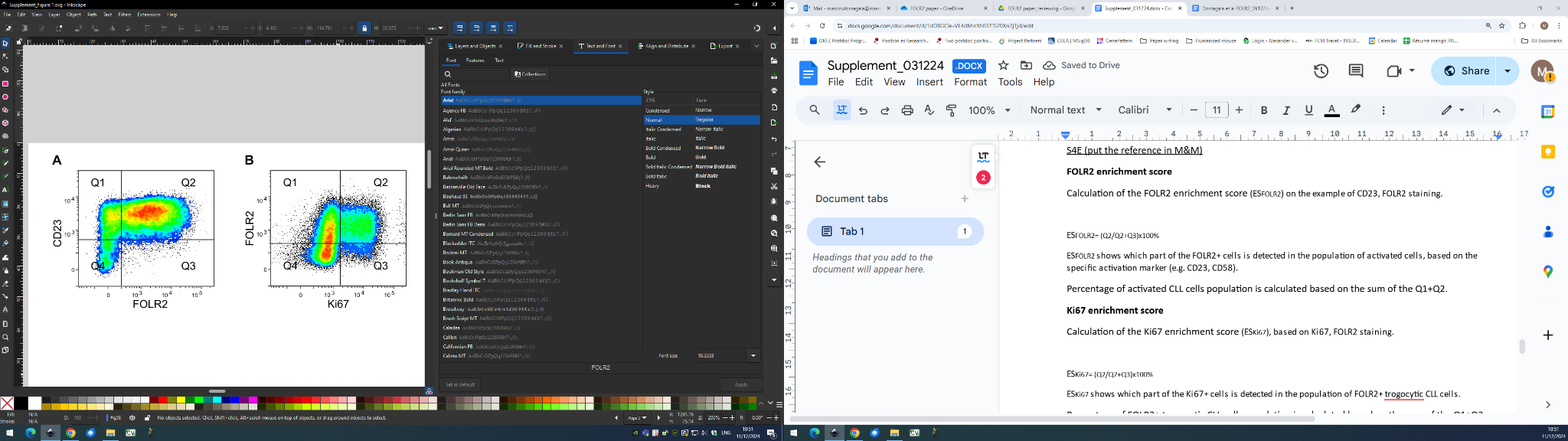

**Figure S4F. Calculation of enrichment scores for evaluation of relation between CLL cell activation or proliferation status, and FOLR2 acquisition.**

A) Calculation of the FOLR2 enrichment scores (ES_FOLR2_) on the example of CD23/FOLR2 staining:

ES_FOLR2_= (Q2/Q2+Q3) × 100%

ES_FOLR2_ indicates a proportion of the FOLR2^+^ cells (Q2) detected in the subpopulation of activated cells (Q1+Q2). Activated CLL cells defined by expression of a specific activation marker (e.g., CD23, CD58).

B) Calculation of the Ki67 enrichment score (ES_Ki67_):

ES_Ki67_= (Q2/Q2+Q3) × 100%

ES_Ki67_ indicates a proportion of Ki67^+^ cells (Q2) detected in the population of FOLR2^+^ trogocytic CLL cells (Q1+Q2).

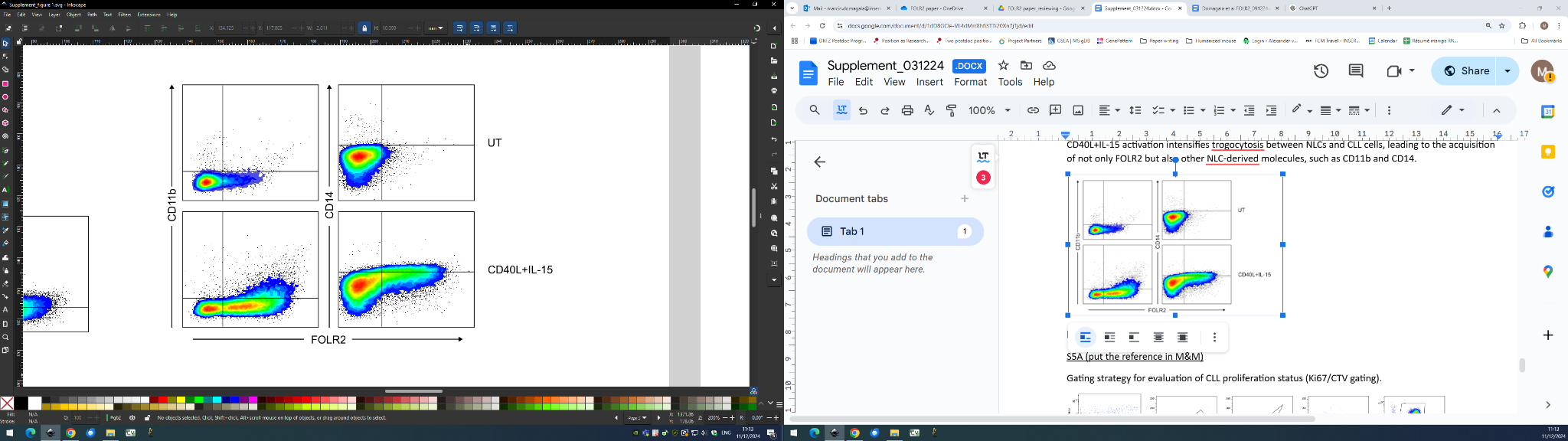

**Figure S4G. CD40L+IL-15 activation intensifies trogocytosis between NLCs and CLL cells**. Representative dot plots from flow cytometry analysis of FOLR2, CD11b and CD14 signals on CLL cells following co-culture with HD-NLCs, depending on CD40L+IL-15 activation. UT – untreated cells.

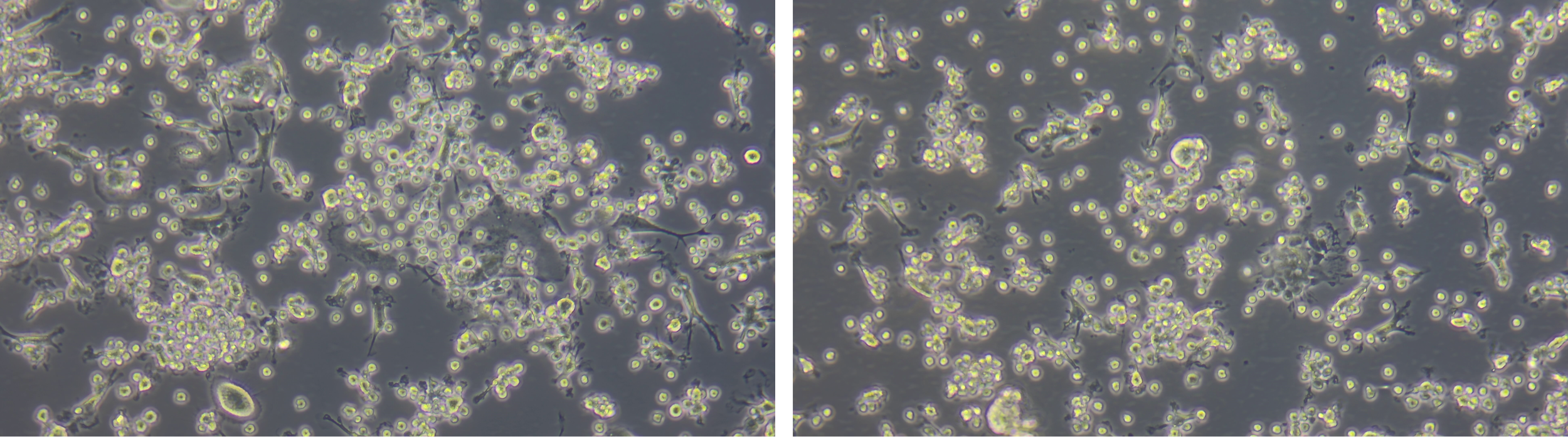

**Figure S4E. Close contact between CLL cells and HD-NLC following CD40L+IL15 stimulation.** HD-NLC-CLL cell co-cultures were activated with CD40L and IL-15. 96h later, floating cells were gently removed by two PBS washes to expose CLL cells bound to NLCs, which were then imaged using bright-field microscopy.

**Supplementary Figure S5**

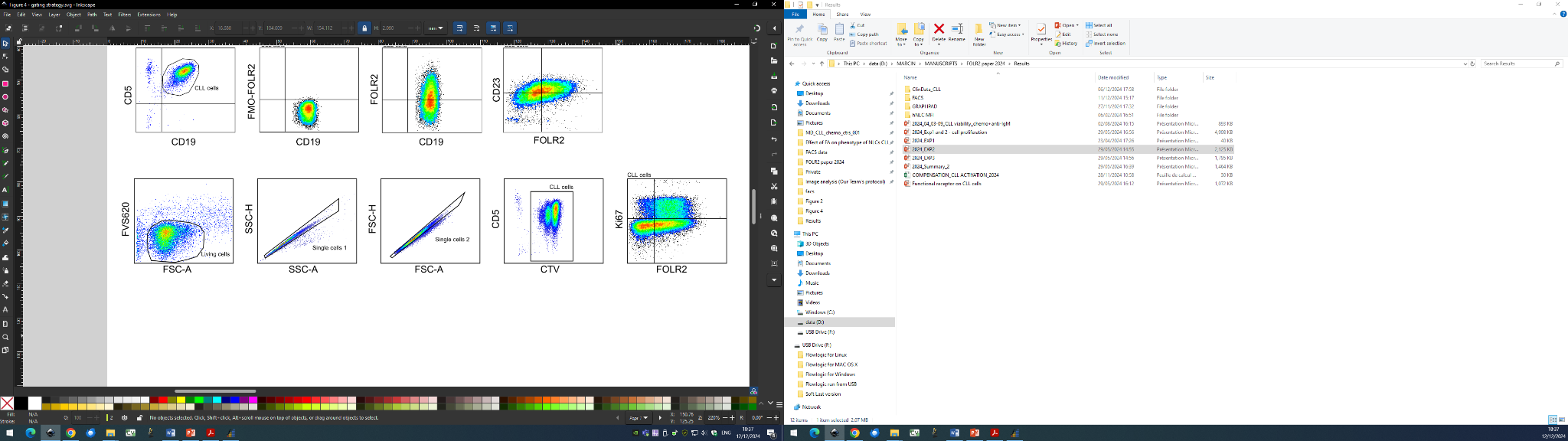

**Figure S5A. Gating strategy for evaluation of CLL proliferation and FOLR2 acquisition.** Dead cells were identified based on positive staining with Fixable Viability Stain 620 (FVS620) and forward scatter (FSC). Cell doublets were excluded based on FSC and side-scatter (SSC) parameters. CLL cells were gated based on positive signal of CTV and staining with anti-CD5 antibody. Positive gate for the FOLR2 and Ki67 signal was determined with isotype control antibodies.

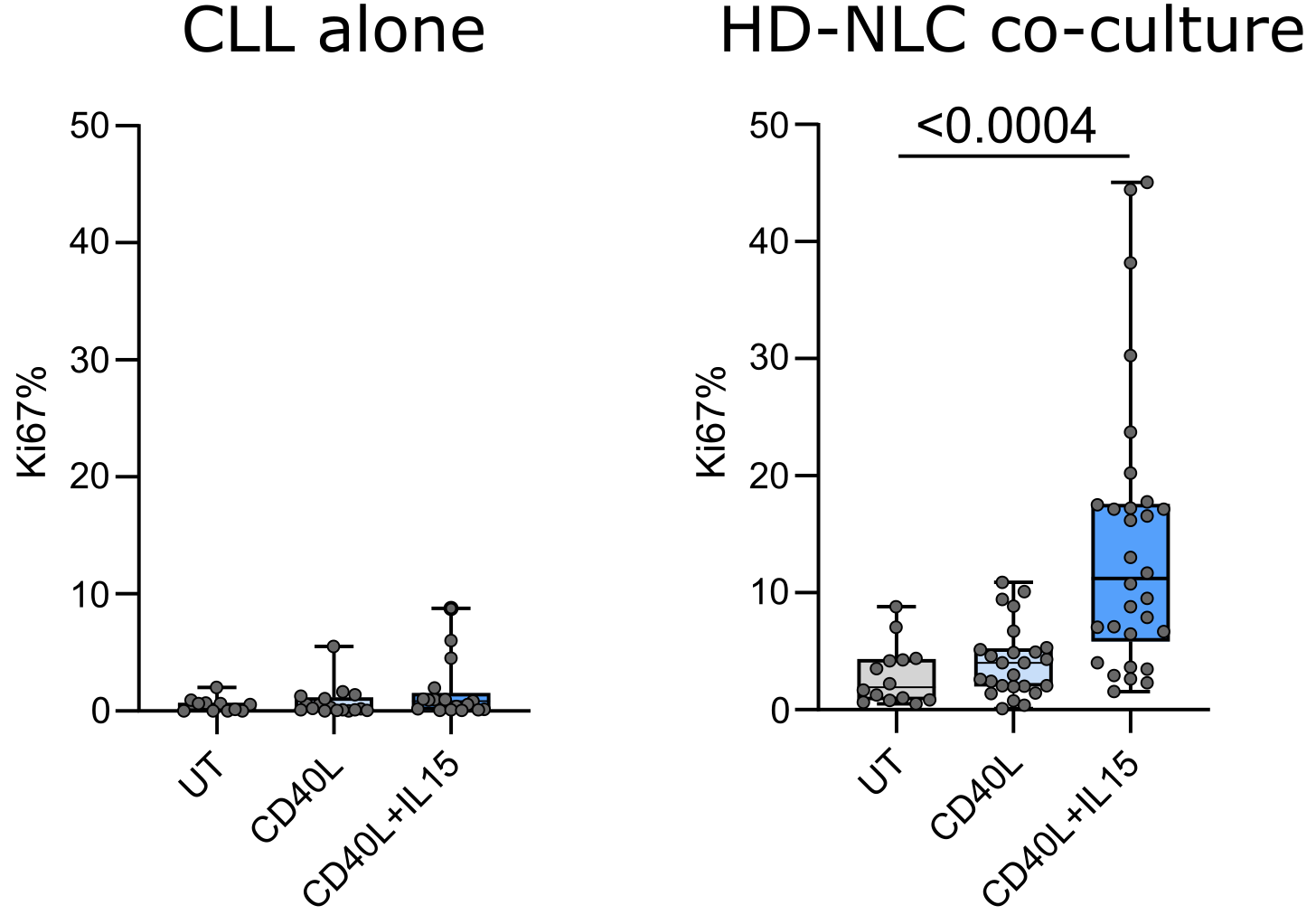

**Figure S5B. IL-15 in combination with CD40L induces robust proliferation of CLL cells in the presence of HD-NLCs.** Effect of CD40L (light blue) and CD40L+IL-15 (blue) activation on Ki67 expression by CLL cells cultivated alone (left) or with HD-NLCs (right) was evaluated with flow cytometry. The box-and-whisker plots display data from n = 4 independent experiments, CLL cells from 9 patients, and HD-NLCs generated from monocytes from 7 healthy donors. Repeated-measures ANOVA with Geisser-Greenhouse correction was used for data analysis, with p-values < 0.05 considered statistically significant.

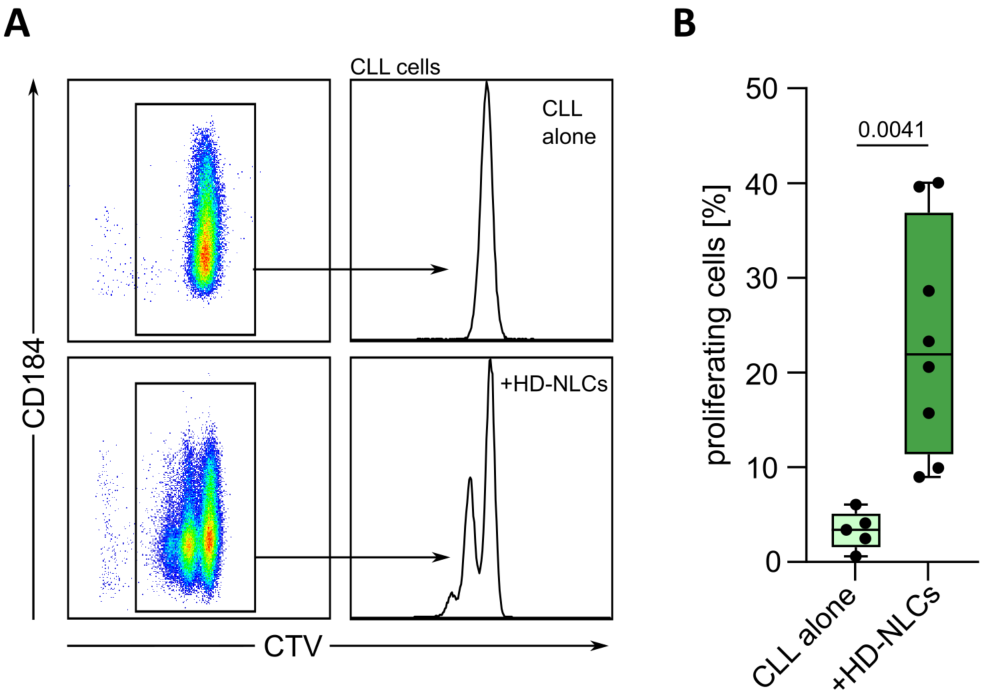

**Figure S5C. CD40L+IL-15 activation leads to increased CLL proliferation in the presence of HD-NLC.** Cancer cell proliferation was measured for monocultures (CLL alone) or co-cultures with HD-NLCs, 5 days after CD40L+IL-15 treatment. A) Representative flow cytometry results. B) Cumulative data from n = 8 patient samples showing the percentage of proliferating CLL cells based on CellTrace™ Violet (CTV) signal reduction. Results were evaluated with paired, two-tailed t-test, with p-values < 0.05 considered significant.

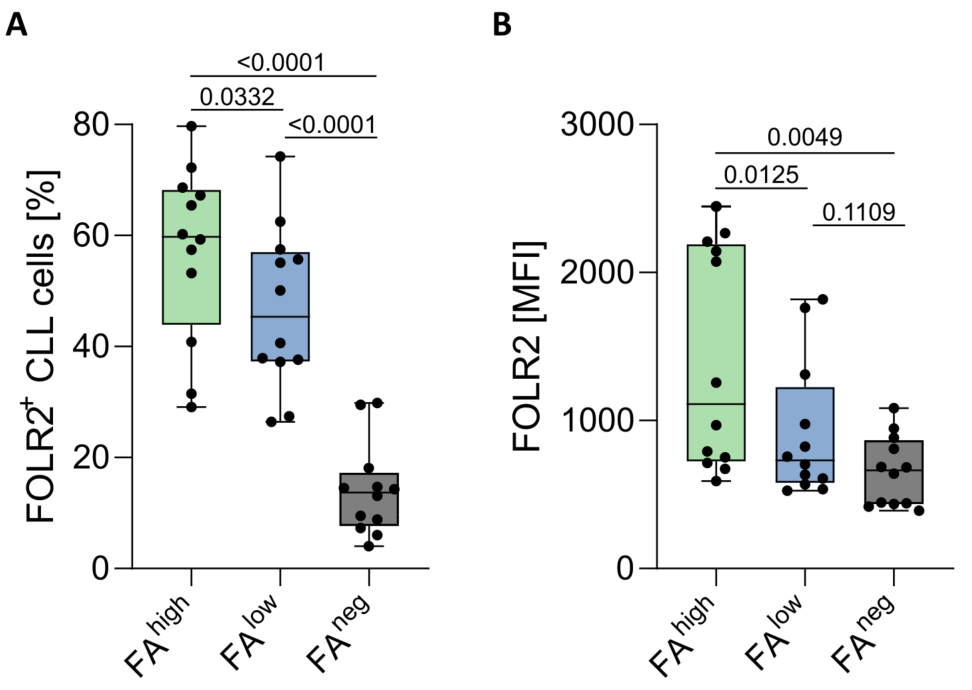

**Figure S5D. Impact of folic acid availability on FOLR2 acquisition by CLL cells.** CLL cells were co-cultured with HD-NLCs in the media with decreasing content of folic acid (FA): FA^high^, FA^low^ and FA^new^, and analyzed by flow cytometry (n = 2 independent experiments; 8 different patient samples; 12 measurement points). A) Comparison of FOLR2^+^ CLL cell proportion, depending on FA content. B) Comparison of FOLR2 signals on FOLR2^+^ CLL cell subpopulation, depending on FA content. MFI - median fluorescence intensity. Results were analyzed by repeated-measures ANOVA with Geisser-Greenhouse correction, with p-values < 0.05 considered statistically significant.

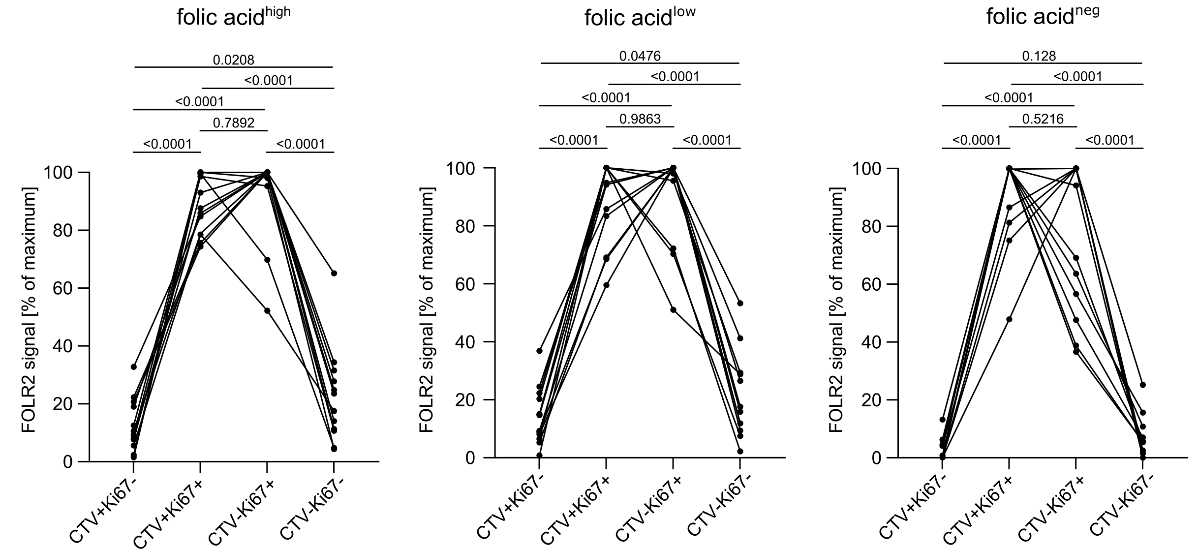

**Figure S5E. FOLR2^high^CLL cells are a subpopulation of actively proliferating cancer cells.** CLL cells were co-cultured with HD-NLCs in the media with decreasing content of folic acid (FA): FA^high^, FA^low^ and FA^new^, and analyzed by flow cytometry to assess FOLR2 levels in relation to proliferative status of CLL cells. CLL cells were divided into four subpopulations (according to Fig. 5E): P1 = CTV^high^/Ki67⁻ (quiescent, non-dividing); P2 = CTV^high^/Ki67⁺ (right before the first division); P3 = CTV^low^/Ki67⁺ (actively cycling, after division); P4 = CTV^low^/Ki67⁻ (post-cycling, quiescent again). To account for heterogeneity in FOLR2 signals, FOLR2 MFI values for each patient sample and experimental condition were normalized to the highest-expressing subpopulation and presented as relative FOLR2 MFI (% of maximum). Repeated-measures ANOVA with Geisser-Greenhouse correction was used to compare FOLR2 expression by different subpopulations of CLL cells depending on their proliferation status (n = 2 independent experiments, 8 patient samples, 12 measurement points), with p-values < 0.05 considered statistically significant.

**Supplementary Figure S6**

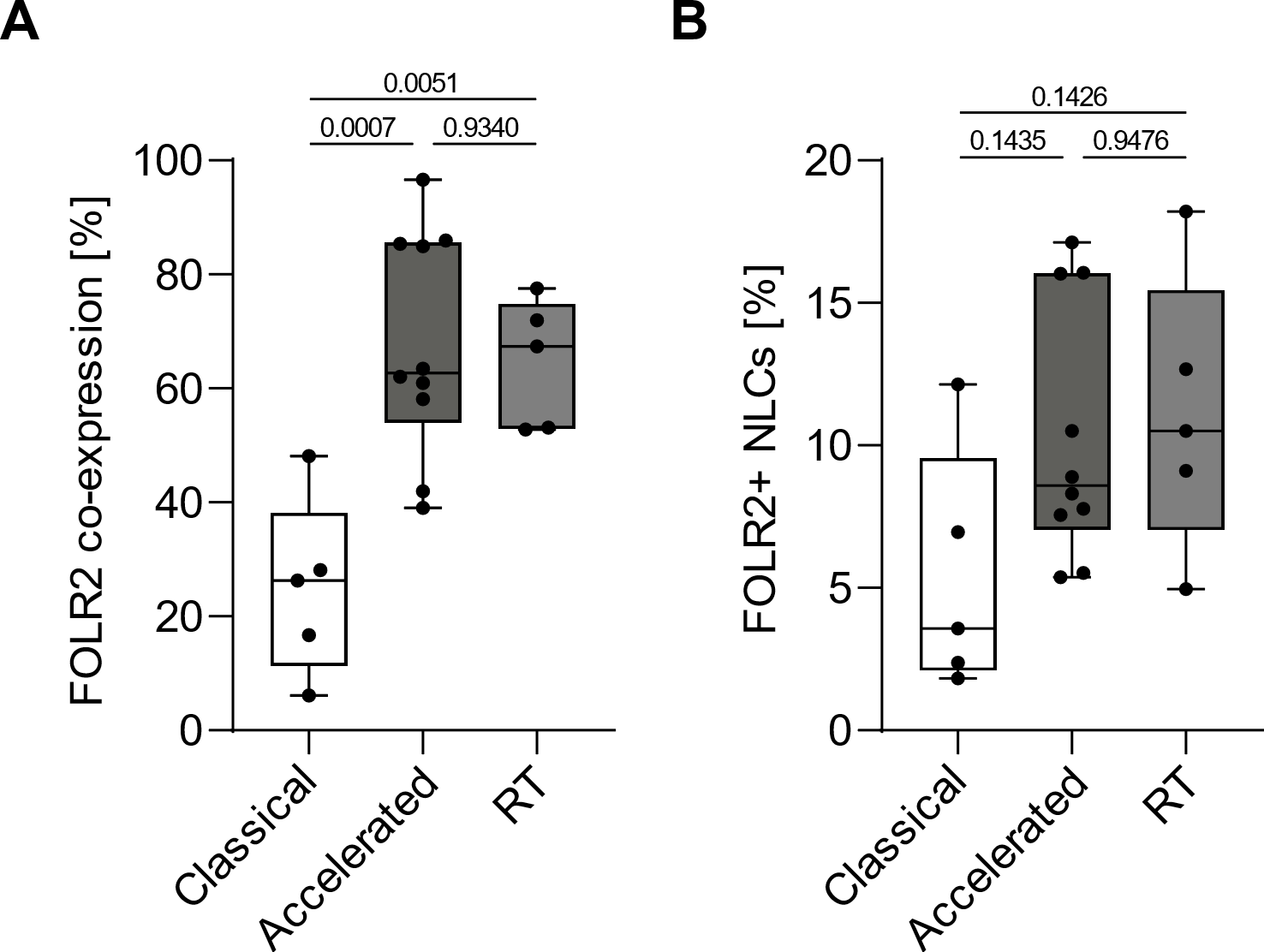

**Figure S6A. Comparison of the frequency of FOLR2⁺ NLCs among CLL subtypes of increasing aggressiveness.** Whole lymph node sections from patients with classical CLL (n = 5), accelerated CLL (n = 10) and CLL with Richter Transformation (RT, n = 5), were analyzed by multiplex immunofluorescence. Differences in the frequency of FOLR2⁺ NLCs among groups were assessed using one-way ANOVA, with P-values < 0.05 considered statistically significant.

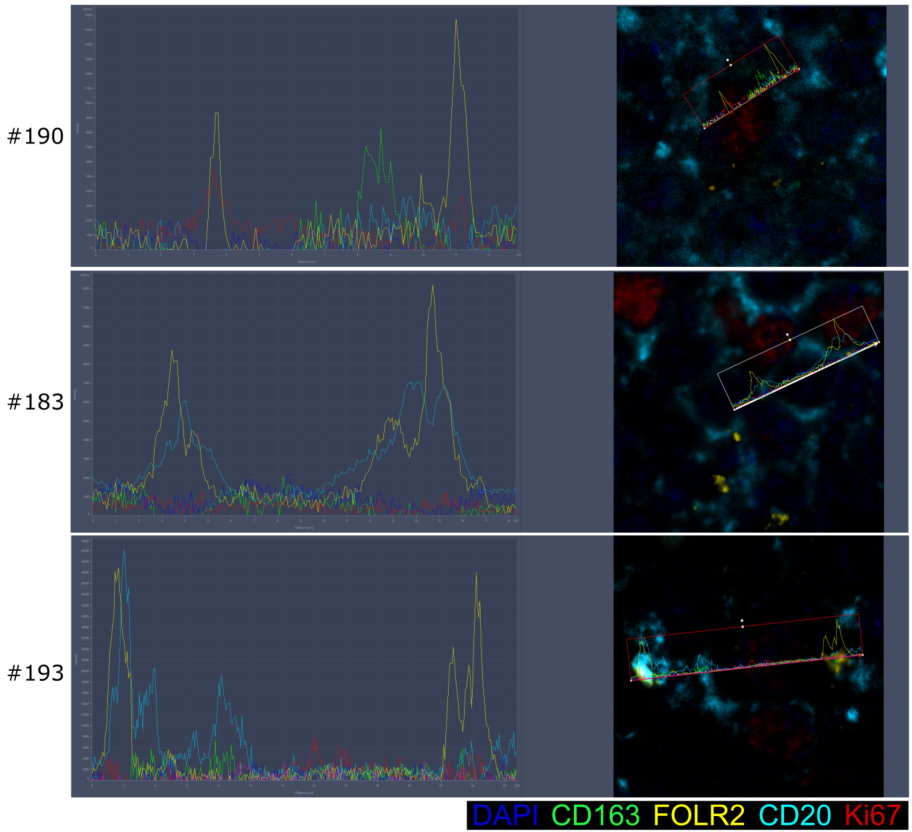

**Figure S6B. Visualization of FOLR2⁺ CLL cells in lymph nodes of patients.** To detect trogocytic cancer B cells *in situ*, lymph nodes from patients #190, #183, and #193 were imaged using a Zeiss 980 confocal microscope (Plan-Apochromat 40×/1.3 oil objective), focusing on regions devoid of NLCs. Subsequently, overlap between FOLR2 (yellow) signals and CD20^+^ (cyan) plasma membrane of cancer cells or their intracellular region, was visualized by co-localization profiles of the selected z-stack sections (left), as indicated with white arrows on the corresponding image (right).
